# Example of Methylome Analysis with MethylIT using Cancer Datasets

**DOI:** 10.1101/261982

**Authors:** Robersy Sanchez, Sally Mackenzie

## Abstract

Methyl-IT, a novel methylome analysis procedure based on information thermodynamics and signal detection was recently released. Methylation analysis involves a signal detection problem, and the method was designed to discriminate methylation regulatory signal from background noise induced by thermal fluctuations. Methyl-IT enhances the resolution of genome methylation behavior to reveal network-associated responses, offering resolution of gene pathway influences not attainable with previous methods. Herein, an example of MethylIT application to the analysis of breast cancer methylomes is presented.

## 1 MethylIT

MethylIT is an R package for methylome analysis based on information thermodynamics and signal detection. The information thermodynamics-based approach is postulated to provide greater sensitivity for resolving true signal from the thermodynamic background within the methylome (Sanchez and Mackenzie 2016). Because the biological signal created within the dynamic methylome environment characteristic of plants is not free from background noise, the approach, designated MethylIT, includes the application of signal detection theory (Greiner, Pfeiffer, and Smith 2000; Carter et al. 2016; Harpaz et al. 2013; Kruspe et al. 2017). A basic requirement for the application of signal detection is a probability distribution of the background noise. Probability distribution, as a Weibull distribution model, can be deduced on a statistical mechanical/thermodynamics basis for DNA methylation induced by thermal fluctuations (Sanchez and Mackenzie 2016). Assuming that this background methylation variation is consistent with a Poisson process, it can be distinguished from variation associated with methylation regulatory machinery, which is non-independent for all genomic regions (Sanchez and Mackenzie 2016). An information-theoretic divergence to express the variation in methylation induced by background thermal fluctuations will follow a Weibull distribution model, provided that it is proportional to minimum energy dissipated per bit of information from methylation change. Herein, we provide an example of MethylIT application to the analysis of breast cancer methylomes. Due to the size of human methylome the current example only covers the analysis of chromosome 13. A full description of MethylIT application of methylome analysis in plants is given in the manuscript (Sanchez et al. 2018).

### 1.1 Installation of MethylIT

To install MethylIT you might need to install the Bioconductor packages: ‘GenomicFeatures’, ‘VariantAnnotation’, ‘ensembldb’, ‘GenomicRanges’, ‘BiocParallel’, ‘biovizBase’, ‘DESeq2’, and ‘genefilter’. Please check that both the R and bioconductor packages are up to date:

`update.packages(ask = FALSE)`

source(‘https://bioconductor.org/biocLite.R’)

`biocLite(ask = FALSE)`

MethylIT can be installed from PSU’s GitLab by typing in an R console:

`install.packages(‘devtools’)`

devtools::install_git(‘https://git.psu.edu/genomath/MethylIT’)

Some possible troubleshooting installation on Ubuntu is given in section S1. Installation on our Windows OS machines was straightforward.

## 2 Available datasets and reading

Methylome datasets of whole-genome bisulfite sequencing (WGBS) are available at Gene Expression Omnibus (GEO DataSets). For the current example, datasets from breast tissues (normal and cancer) and embryonic stem cells will be downloaded from GEO. The data set are downloaded providing the GEO accession numbers for each data set to the function ‘getGEOSuppFiles’ (for details type ?getGEOSuppFiles in the R console).

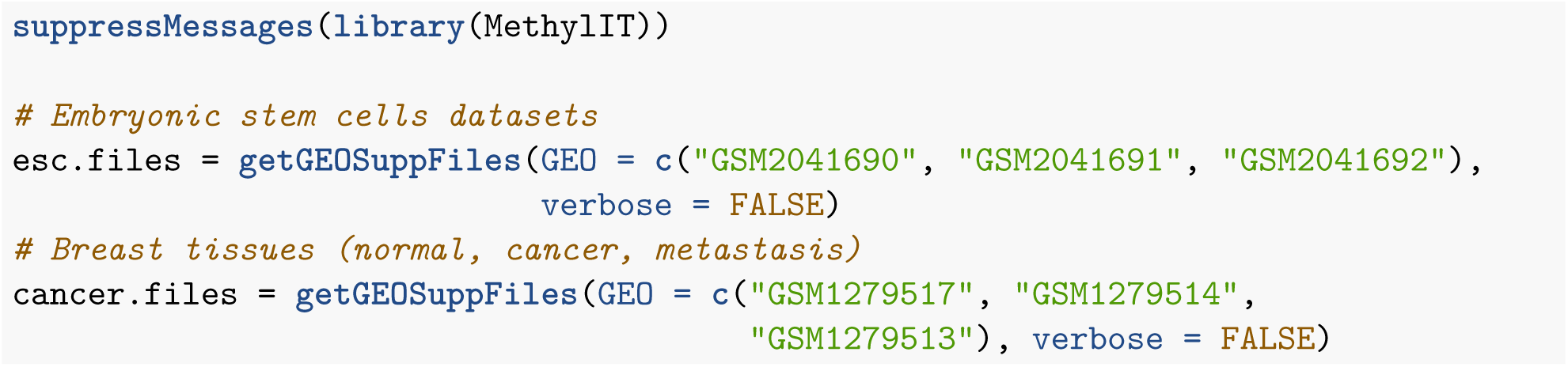

The file path and name of each downloaded dataset is found in the output variables ‘esc.files’ and ‘cancer.files’.

### 2.1 Reading datasets

Datasets for our example can be read with function ‘readCounts2GRangesList’. To specify the reading of only chromosome 13, we can specify the parameter ‘chromosomes = “Chr13”’. The symbol chromosome 13, in this case “Chr13”, must be consistent with the annotation provided in the given GEO dataset. Each file is wholly read with the setting ‘chromosomes = “Chr13”’ and then the GRanges are built only with chromosome 13, which could be time consuming. However, users working on Linux OS can specify the reading of specific lines from each file by using regular expressions. For example, if only chromosomes 1 and 3 are required, then we can set chromosomes = NULL (default) and ‘chromosome.pattern = “ˆChr[1,3]”’. This will read all the lines in the downloaded files starting with the words “Chr1” or “Chr3”. If we are interested in chromosomes 1 and 2, then we can set ‘chromosome.pattern = “ˆChr[1-2]”’. If all the chromosomes are required, then set chromosomes = NULL and chromosome.pattern = NULL (default).

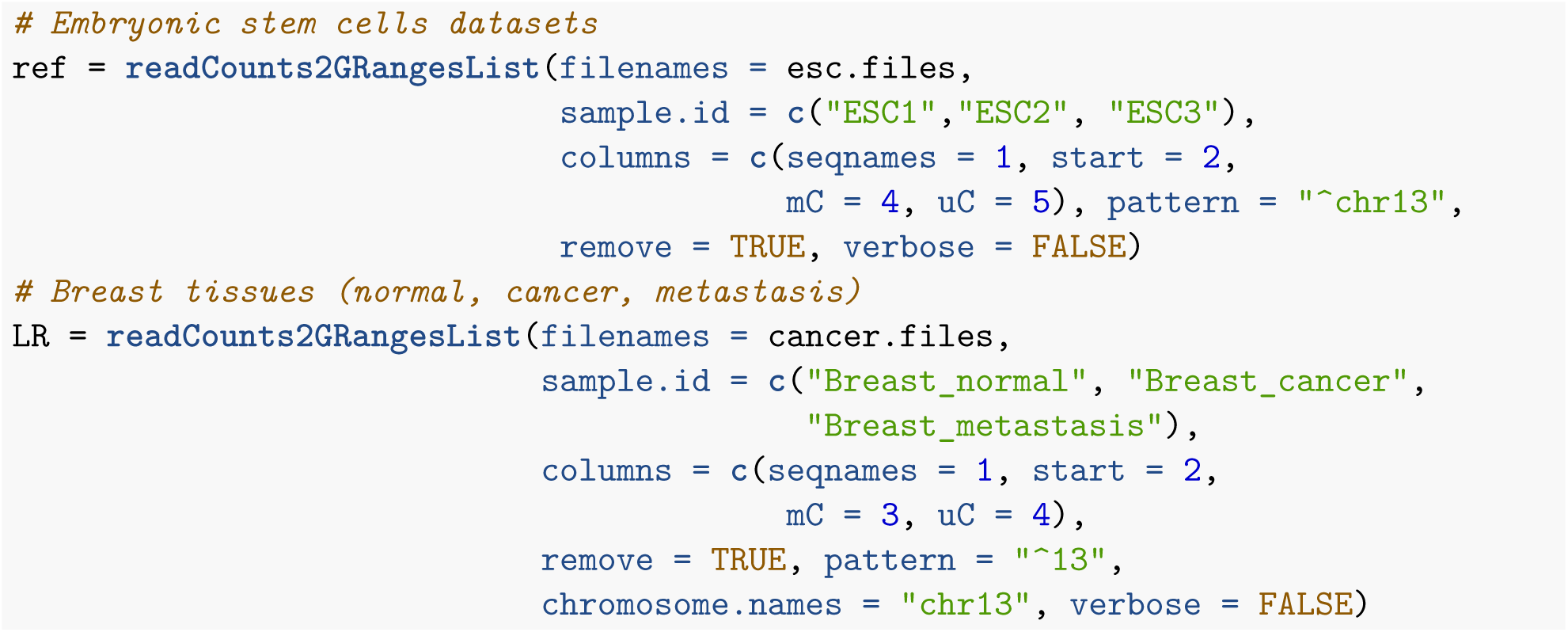

In the metacolumn of the last GRanges object, mC and uC stand for the methylated and unmethylated read counts, respectively. Notice that option ‘remove = TRUE’ remove the decompressed files (default: FALSE, see ?readCounts2GRangesList for more details about this function).

## 3. The reference individual

Any two objects located in a space can be compared if, and only if, there is a reference point (a coordinate system) in the space and a metric. Usually, in our daily 3D experience, our brain automatically sets up the origin of coordinates equal to zero. The differences found in the comparison depend on the reference used to perform the measurements and from the metric system. The space where the objects are located (or the set of objects) together with the metric is called metric space.

To evaluate the methylation differences between individuals from control and treatment we introduce a metric in the bidimensional space of methylation levels *P*_*i*_ = (*p*_*i*_, 1 − *P*_*i*_). Vectors *P*_*i*_ provide a measurement of the uncertainty of methylation levels. However, to perform the comparison between the uncertainty of methylation levels from each group of individuals, control (*c*) and treatment (*t*), we should estimate the uncertainty variation with respect to the same individual reference on the mentioned metric space. The reason to measure the uncertainty variation with respect to the same reference resides in that even sibling individuals follow an independent ontogenetic development. This a consequence of the “omnipresent” action of the second law of thermodynamics in living organisms. In the current example, we will create the reference individual by pooling the methylation counts from the embryonic stem cells.

It should be noticed that the results are sensitive to the reference used. The statistics mean, median, or sum of the read counts at each cytosine site of some control samples can be used to create a virtual reference sample. It is up to the user whether to apply the ‘row sum’, ‘row mean’ or ‘row median’ of methylated and unmethylated read counts at each cytosine site across individuals:

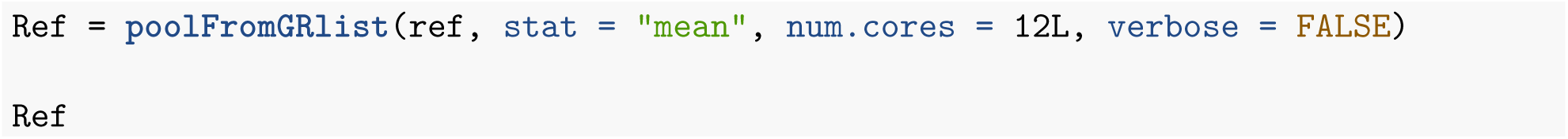

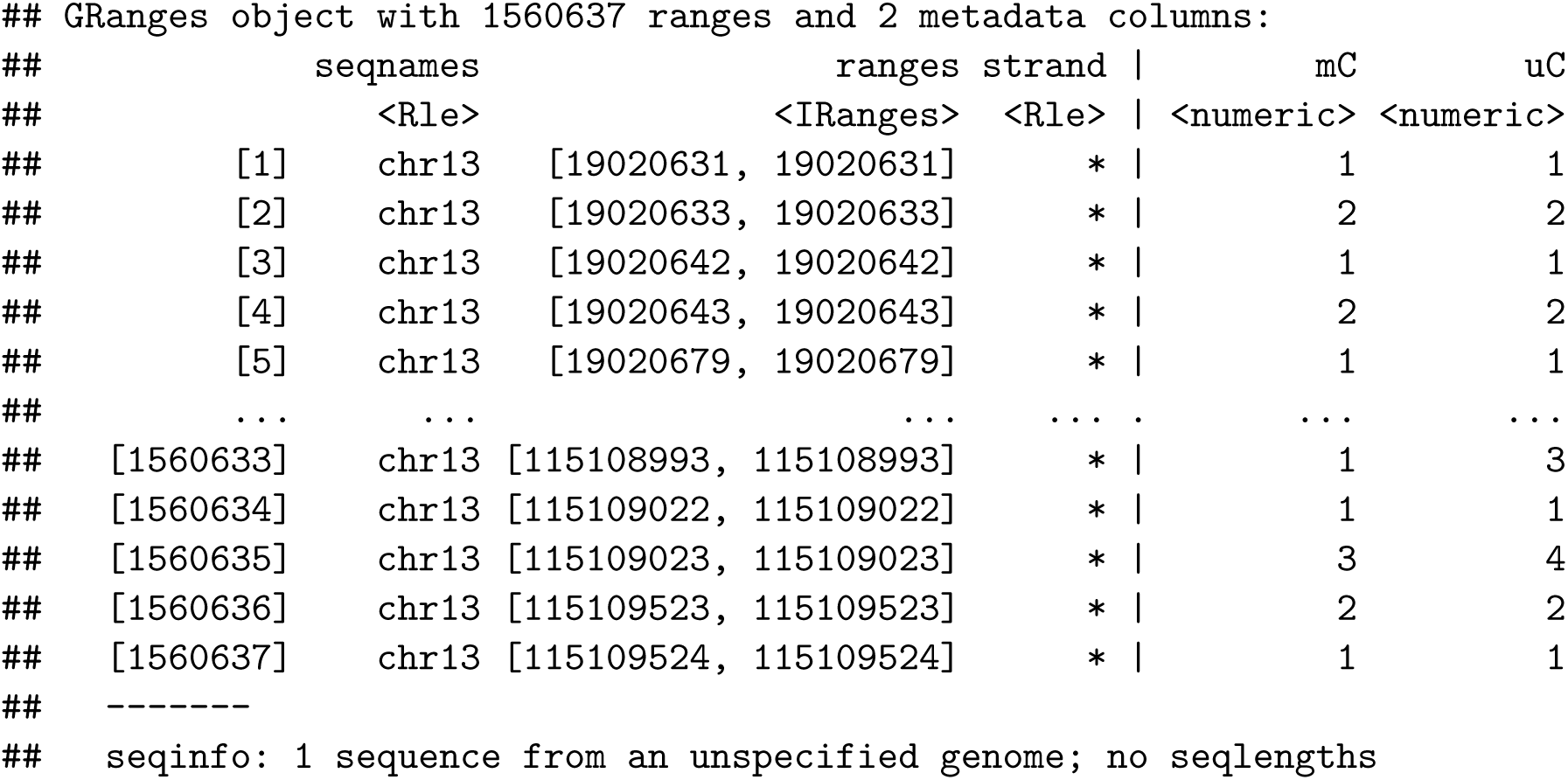

Only direct lab experiments can reveal whether differences detected with distinct references outside the experimental conditions for control and treatment groups are real. The best reference would be estimated using a subset of individuals from control group. Such a reference will contribute to remove the intragroup variation, in control and in treatment groups, induced by environmental changes external to or not controlled by the experimental conditions.

Methylation analysis for each cystosine position is frequently performed in the bidimensional space of (*methylated, unmethylated*) read counts. Frequently, Fisher test is applied to a single cytosine position, under the null hypothesis that the proportions *p*_*ct*_ = *methylated*_*ct*_/(*methylated*_*ct*_ + *unmethylated*_*ct*_) and *p*_*tt*_ = *methylated*_*tt*_/(*methylated*_*tt*_ + *unmethylated*_*tt*_) are the same for control and treatment, respectively. In this case, the implicit reference point for the counts at every cytosine positions is (*methylated* = 0, *unmethylated* = 0), which corresponds to the point *P*_*i*_ = (0, 1).

In our case, the Hellinger divergence (the metric used, here) of each individual in respect to the reference is the variable to test in place of (*methylated, unmethylated*) read counts or the methylation levels *P*_*i*_ = (*p*_*i*_, 1 − *p*_*i*_).

The use of references is restricted by the thermodynamics basis of the the theory. The current information-thermodynamics based approach is supported on the following postulate:

*“High changes of Hellinger divergences are less frequent than low changes, provided that the divergence is proportional to the amount of energy required to process one bit of information in methylation system”*.

The last postulate acknowledges the action of the second law of thermodynamics on the biomolecular methylation system. For the methylation system, it implies that the frequencies of the information divergences between methylation levels must be proportional to a Boltzmann factor (see supplementary information from reference (Sanchez and Mackenzie 2016)). In other words, the frequencies of information divergences values should follow a trend proportional to an exponential decay. If we do not observe such a behaviour, then either the reference is too far from experimental condition or we are dealing with an extreme situation where the methylation machinery in the cell is dysfunctional. The last situation is found, for example, in the silencing mutation at the gene of cytosine-DNA-methyltransferase in Arabidopsis *thaliana*. Methylation of 5-methylcytosine at CpG dinucleotides is maintained by MET1 in plants.

In our current example, the embryonic stem cells reference is far from the breast tissue samples and this could affect the nonlinear fit to a Weibull distribution (see below). To illustrate the effect of the reference on the analysis, a new reference will be built by setting:

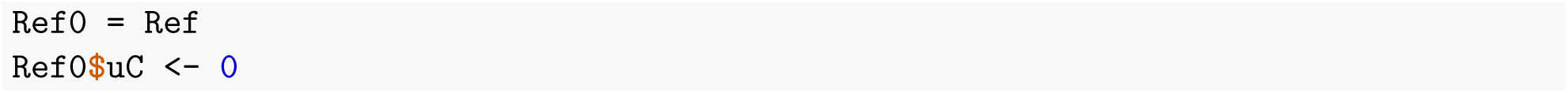

The reason for the above replacement is that natural methylation changes (Ref$mC) obey the second law of thermodynamics, and we do not want to arbitrarily change the number of methylated read counts. ‘mC’ carries information linked to the amount of energy expended in the tissue associated with concrete methylation changes. However, ‘uC’ is not linked to any energy expended by the methylation machinery in the cells. In the bidimensional space *P*_*i*_ = (*p*_*i*_, 1 − *P*_*i*_), reference *Ref0* corresponds to the point *P*_*i*_ = (1, 0) at each cytosine site *i*, i.e., the value of methylation level at every cytosine site in reference *Ref0* is 1. The analyses with respect to both individual references, *Ref* and *Ref0*, will be performed in the downstream steps.

## 4 Hellinger divergence estimation

To perform the comparison between the uncertainty of methylation levels from each group of individuals, control (*c*) and treatment (*t*), the divergence between the methylation levels of each individual is estimated with respect to the same reference on the metric space formed by the vector set *P*_*i*_ = (*p*_*i*_, 1 − *P*_*i*_) and the Hellinger divergence *H*. Basically, the information divergence between the methylation levels of an individual *j* and reference sample *r* is estimated according to the Hellinger divergence given by the formula:

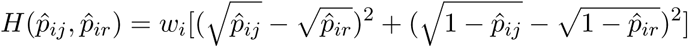

where 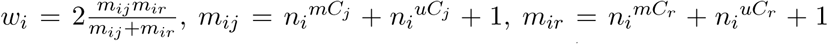 and *j* ϵ {*c, t*}. This equation for Hellinger divergence is given in reference (Basu, Mandal, and Pardo 2010), but other information theoretical divergences can be used as well. Next, the information divergence for control (Breast_normal) and treatment (Breast_cancer and Breast_metastasis) samples are estimated with respect to the reference virtual individual. A Bayesian correction of counts can be selected or not. In a Bayesian framework, methylated read counts are modeled by a beta-binomial distribution, which accounts for both the biological and sampling variations (Hebestreit, Dugas, and Klein 2013; Robinson et al. 2014; Dolzhenko and Smith 2014). In our case we adopted the Bayesian approach suggested in reference (Baldi and Brunak 2001) (Chapter 3). In a Bayesian framework with uniform priors, the methylation level can be defined as: *p* = (*mC* + 1)*/*(*mC* + *uC* + 2).

However, the most natural statistical model for replicated BS-seq DNA methylation measurements is beta-binomial (the beta distribution is a prior conjugate of binomial distribution). We consider the parameter *p* (methylation level) in the binomial distribution as randomly drawn from a beta distribution. The hyper-parameters α and β from the beta-binomial distribution are interpreted as pseudo-counts. The information divergence is estimated here using the function ‘estimateDivergence’:

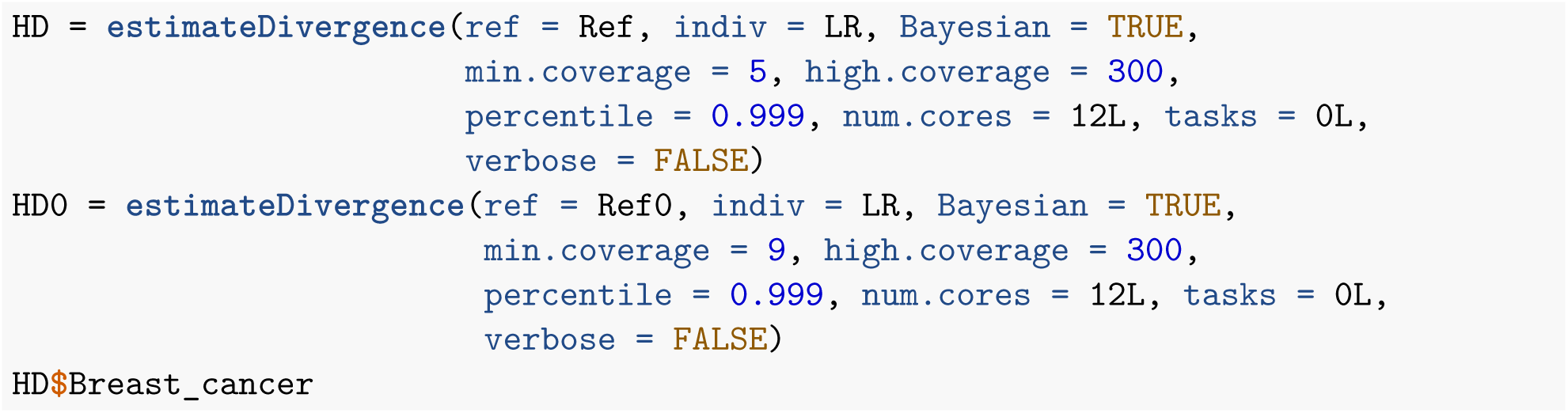

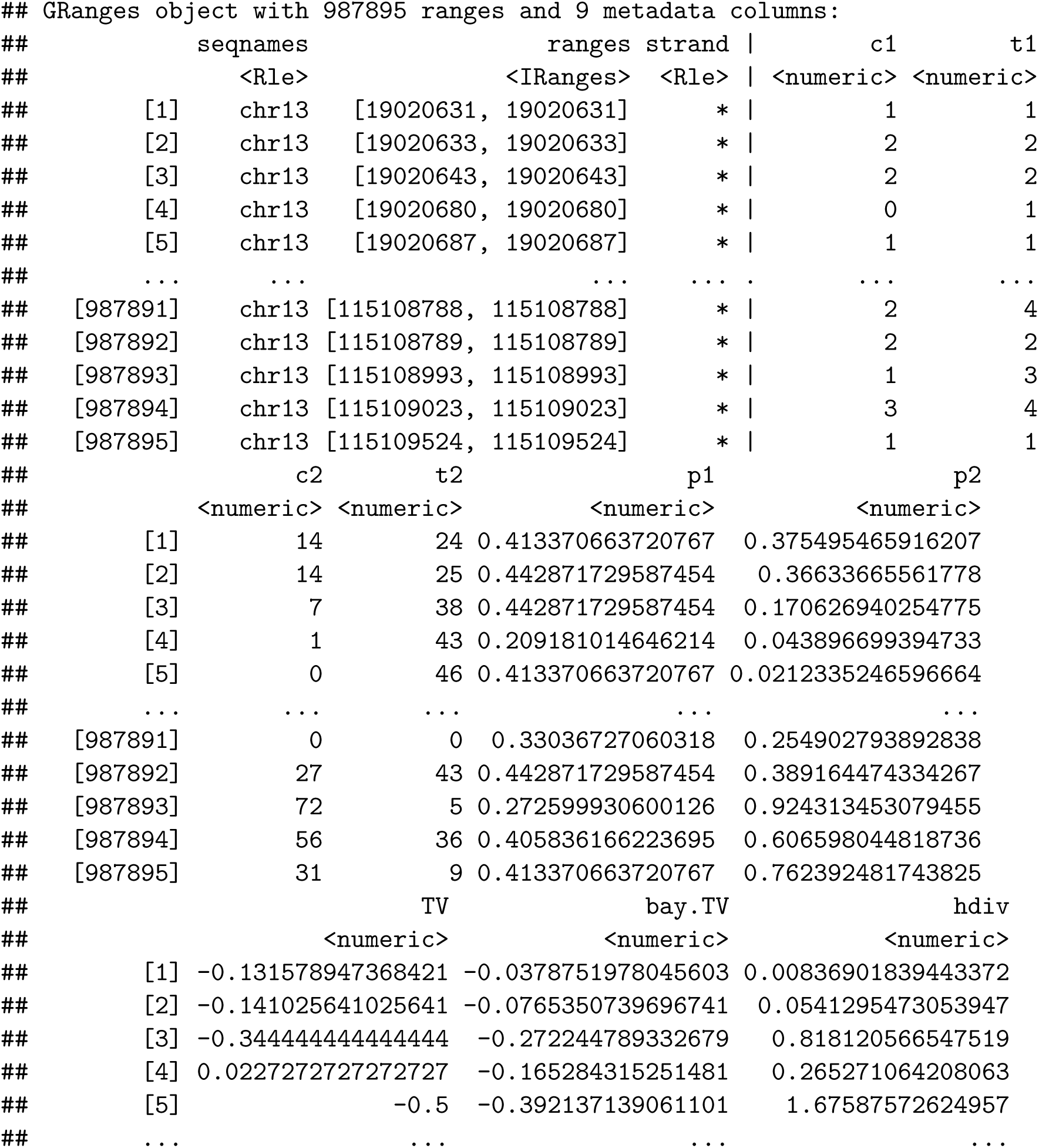
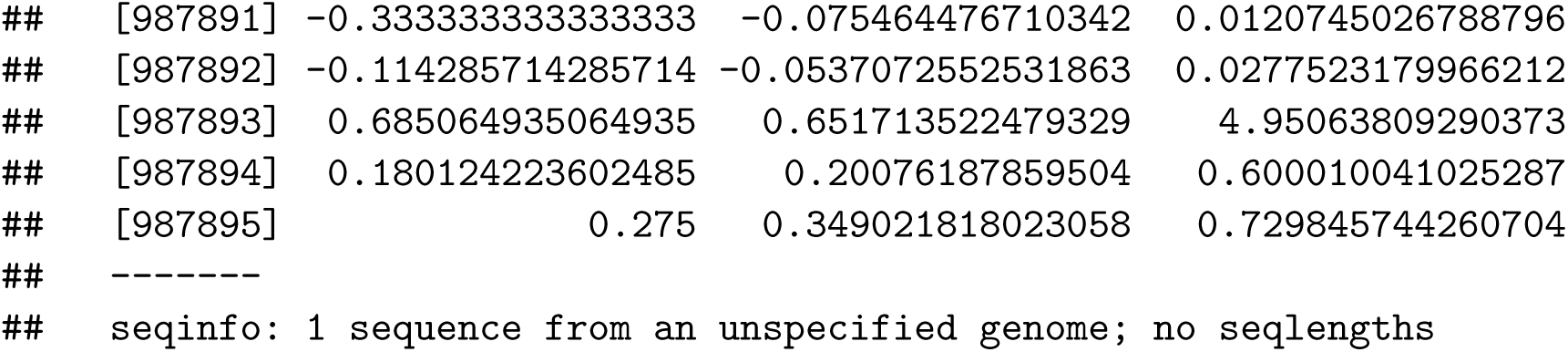

Function ‘estimateDivergence’ returns a list of GRanges objects with the four columns of counts, the information divergence, and additional columns:

1. The original matrix of methylated (*c_i_*) and unmethylated (*t_i_*) read counts from control (*i* = 1) and treatment (*i* = 2) samples.
2. “p1” and “p2”: methylation levels for control and treatment, respectively.
3. “bay.TV”: total variation TV = p2 - p1.
4. “TV”: total variation based on simple counts: *T V* = *c*1*/*(*c*1 + *t*1)*c*2*/*(*c*2 + *t*2).
5. “hdiv”: Hellinger divergence.

If Bayesian = TRUE, results are based on the posterior estimations of methylation levels *p*1 and *p*2. Filtering by coverage is provided at this step which would be used unless previous filtering by coverage had been applied. This is a pairwise filtering. Cytosine sites with ‘coverage’ > ‘min.coverage’ and ‘coverage’ < ‘percentile’ (e.g., 99.9 coverage percentile) in at least one of the samples are preserved. The coverage percentile used is the maximum estimated from both samples: reference and individual.

For some GEO datasets only the methylation levels for each cytosine site are provided. In this case, Hellinger divergence can be estimated as given in reference (Sanchez and Mackenzie 2016):

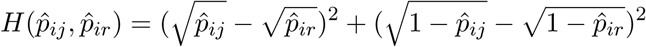

### 4.1. Histogram and boxplots of divergences estimated in each sample

First, the data of interest (Hellinger divergences, “hdiv”) are selected from the GRanges objects:

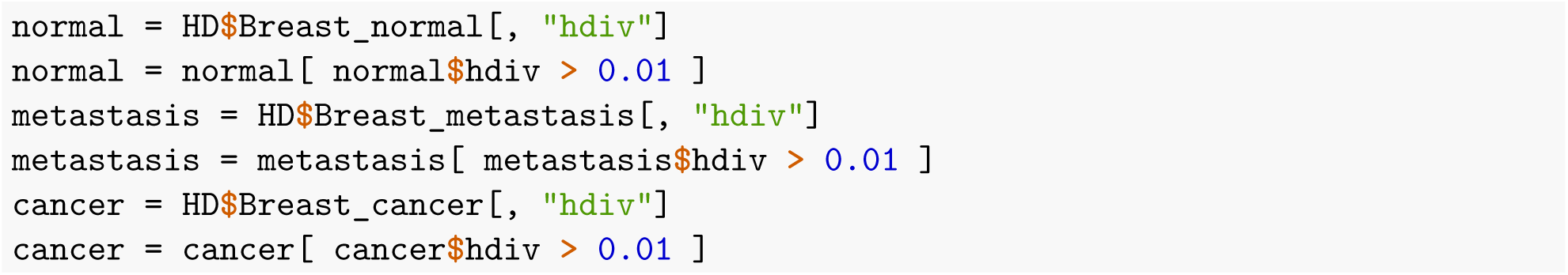

Next, a single GRanges object is built from the above set of GRanges objects using the function ‘uniqueGRanges’. Notice that the number of cores to use for parallel computation can be specified.

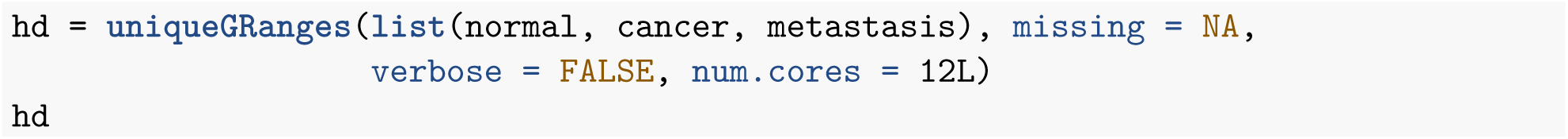

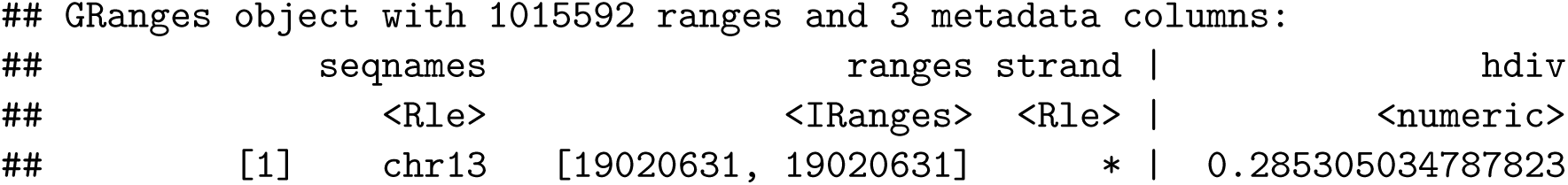
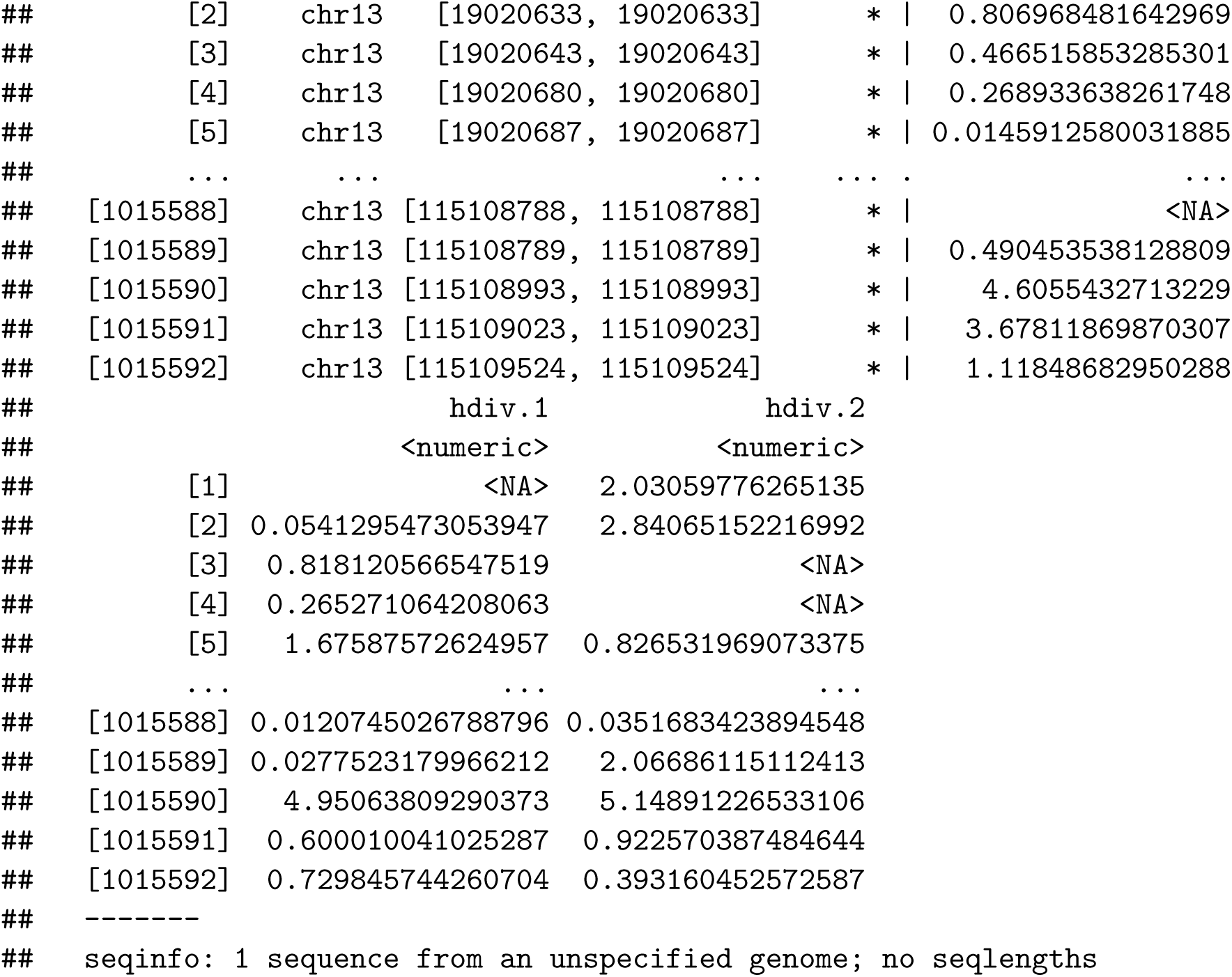

Now, the Hellinger divergences estimated for each sample are in a single matrix on the metacolumn of the GRanges object and we can proceed to build the histogram and boxplot graphics for these data.

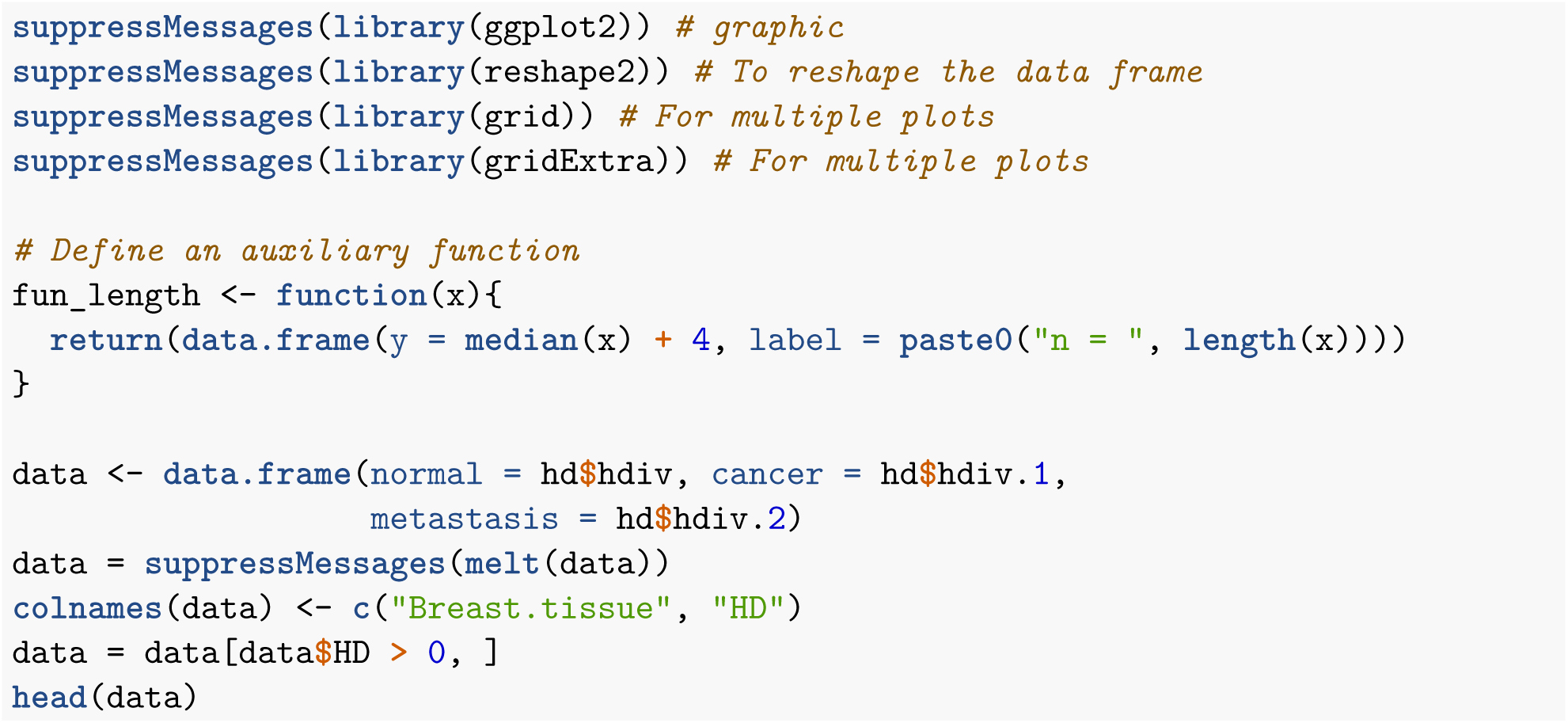

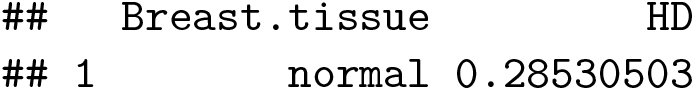
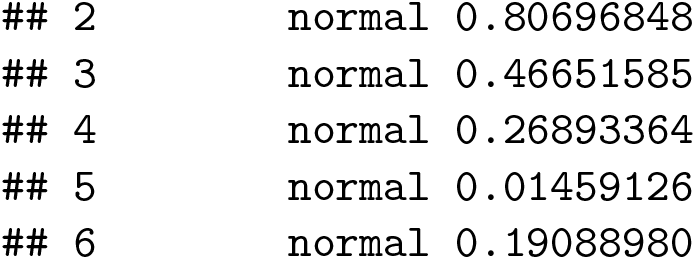

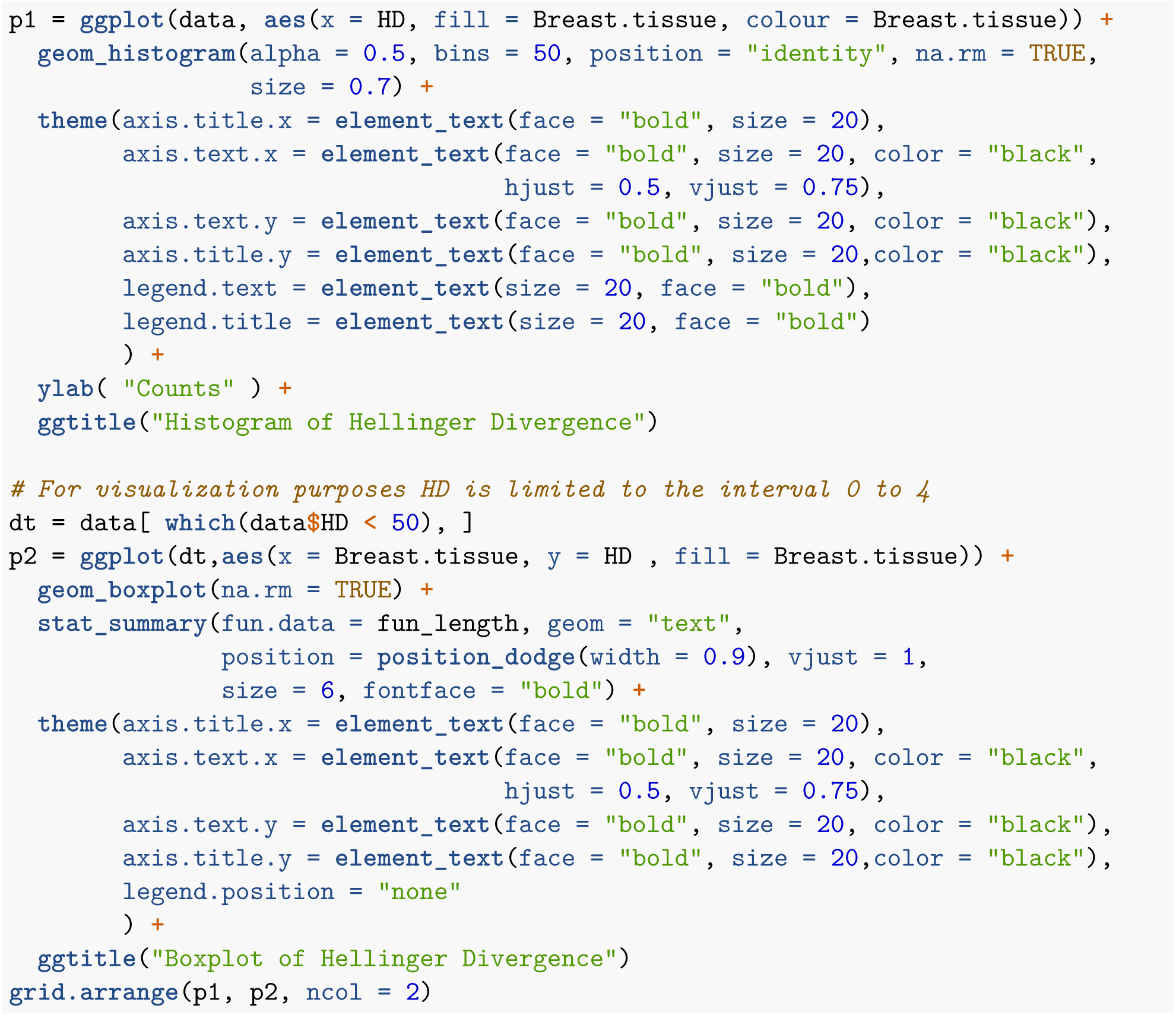

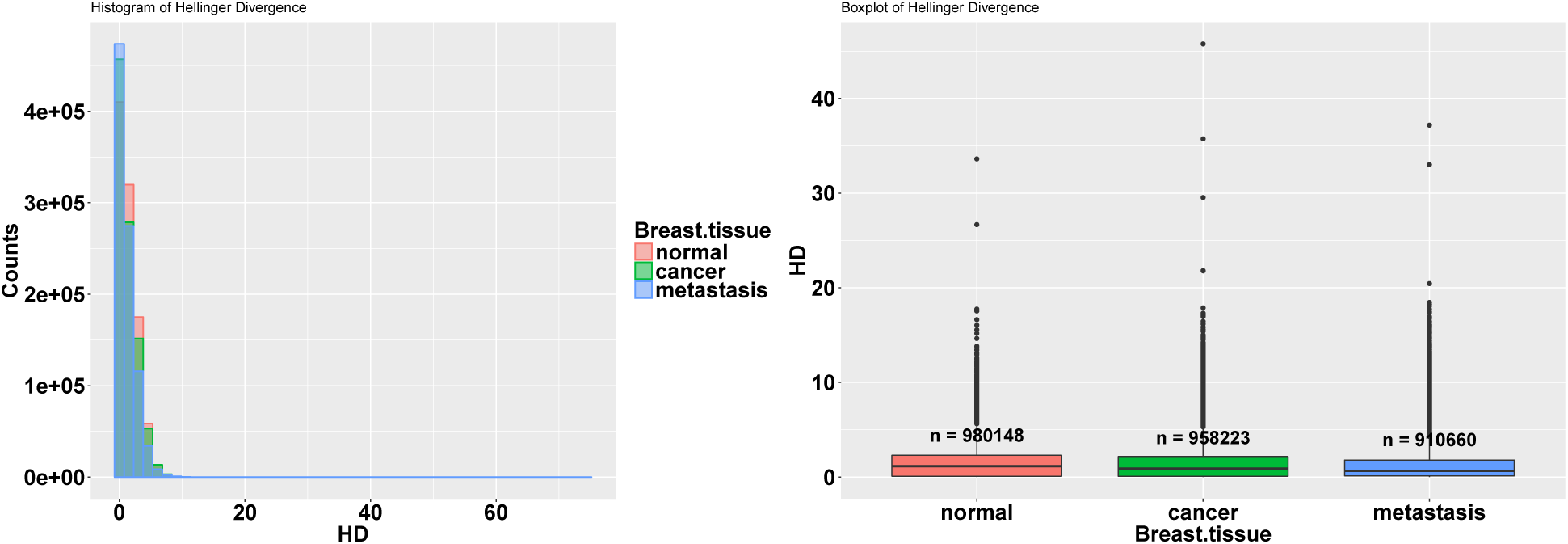

Except for the tail, most of the methylation changes occurred under the area covered by the density curve corresponding to the normal breast tissue. This is theoretically expected. This area is explainable in statistical physical terms and, theoretically, it should fit a Weibull distribution. The tails regions cover the methylation changes that, with high probability, are not induced by thermal fluctuation and are not addressed to stabilize the DNA molecule. These changes are methylation signal. Professor David J. Miller (Department of Electrical Engineering, Penn State) proposed modeling the distribution as a mixed Weibull distribution to simultaneously describe the background methylation noise and the methylation signal (personal communication, January, 2018). This model approach seems to be supported by the above histogram, but it must be studied before being incorporated in a future version of Methyl-IT.

### 5. Nonlinear fit of Weibull distribution

A basic requirement for the application of signal detection is the knowledge of the probability distribution of the background noise. Probability distribution, as a Weibull distribution model, can be deduced on a statistical mechanical/thermodynamics basis for DNA methylation induced by thermal fluctuations (Sanchez and Mackenzie 2016). Assuming that this background methylation variation is consistent with a Poisson process, it can be distinguished from variation associated with methylation regulatory machinery, which is non-independent for all genomic regions (Sanchez and Mackenzie 2016). An information-theoretic divergence to express the variation in methylation induced by background thermal fluctuations will follow a Weibull distribution model, provided that it is proportional to the minimum energy dissipated per bit of information associated with the methylation change. The nonlinear fit to a Weibull distribution model is performed by the function ‘nonlinearFitDist’.

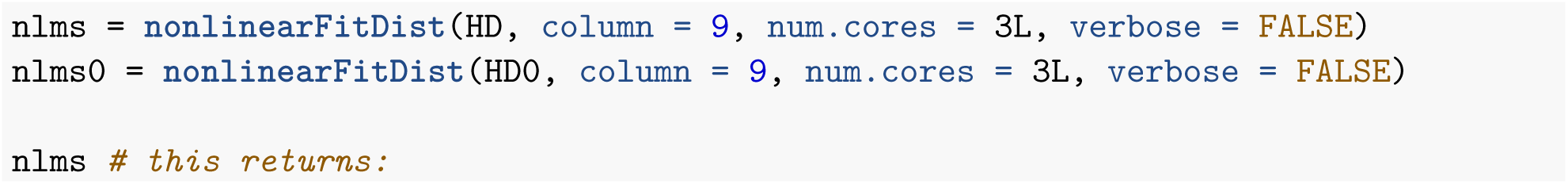

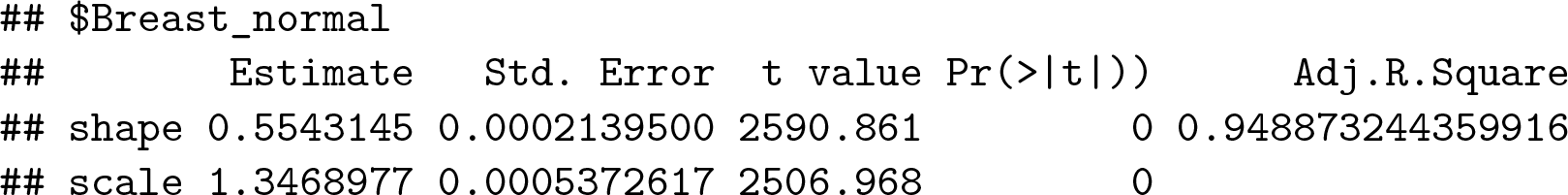
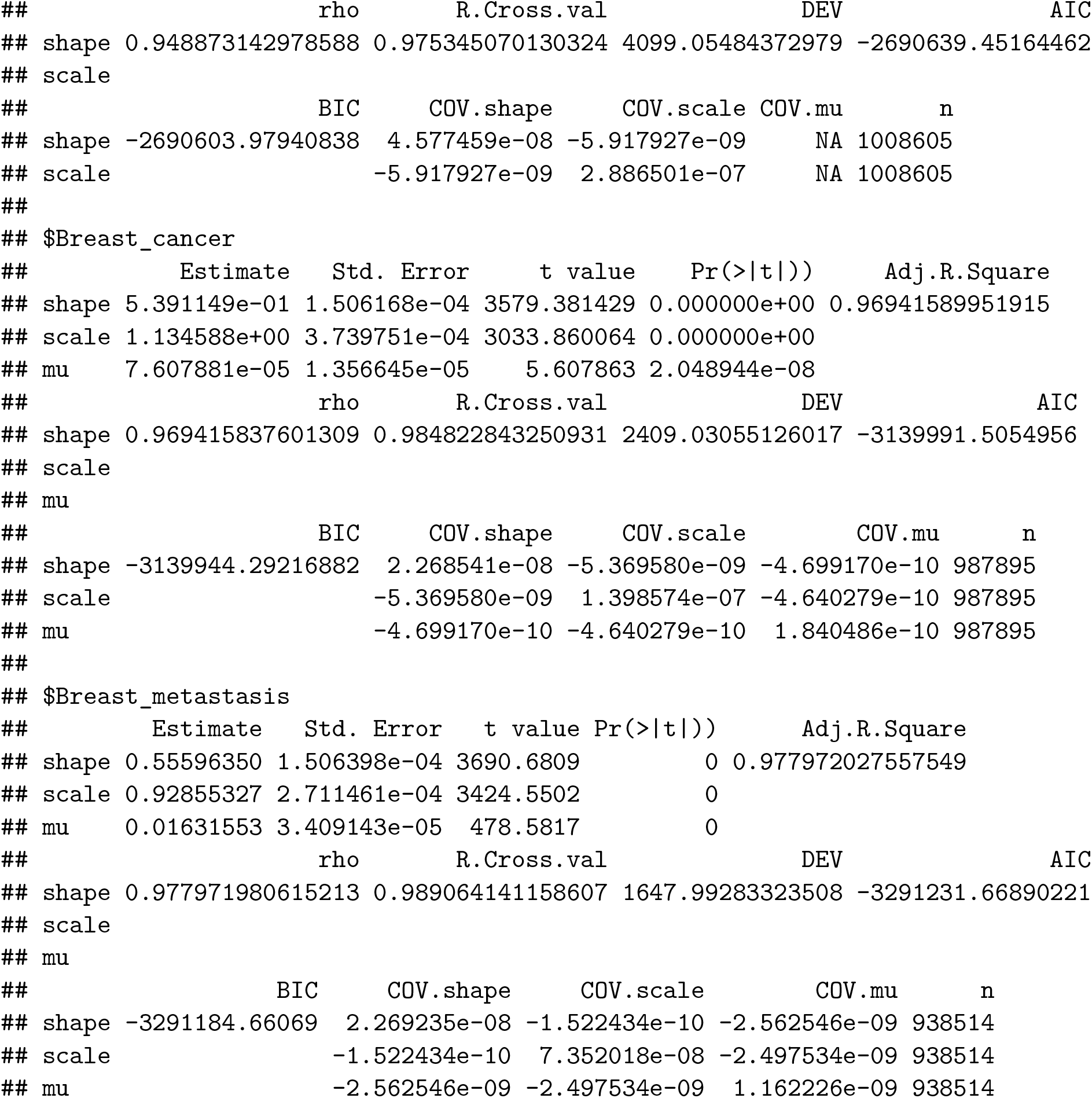

Cross-validations for the nonlinear regressions (R.Cross.val) were performed as described in reference (Stevens 2009). In addition, Stein’s formula for adjusted R squared (ρ) was used as an estimator of the average cross-validation predictive power (Stevens 2009).

The goodness-of-fit of Weibull to the HD0 (*Ref0*) data is better than to HD (*Ref*): `nlms0`

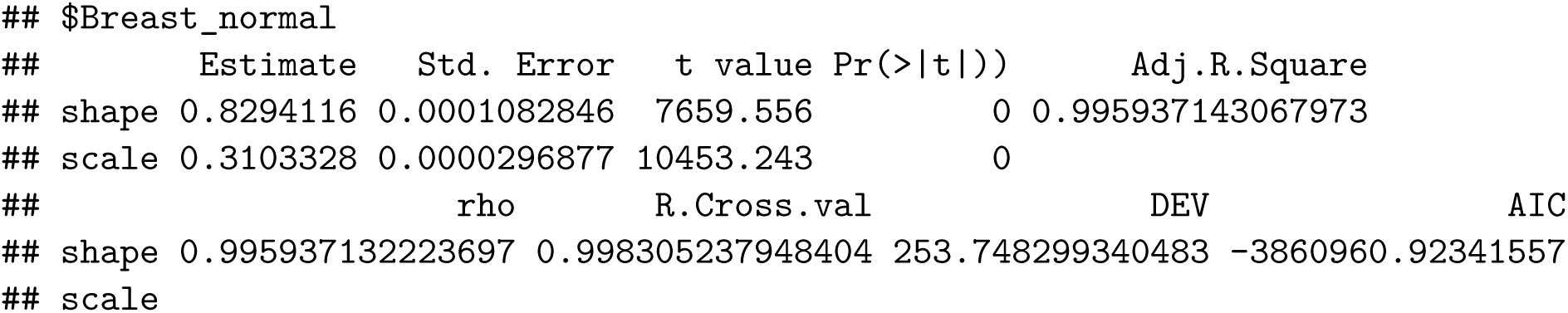
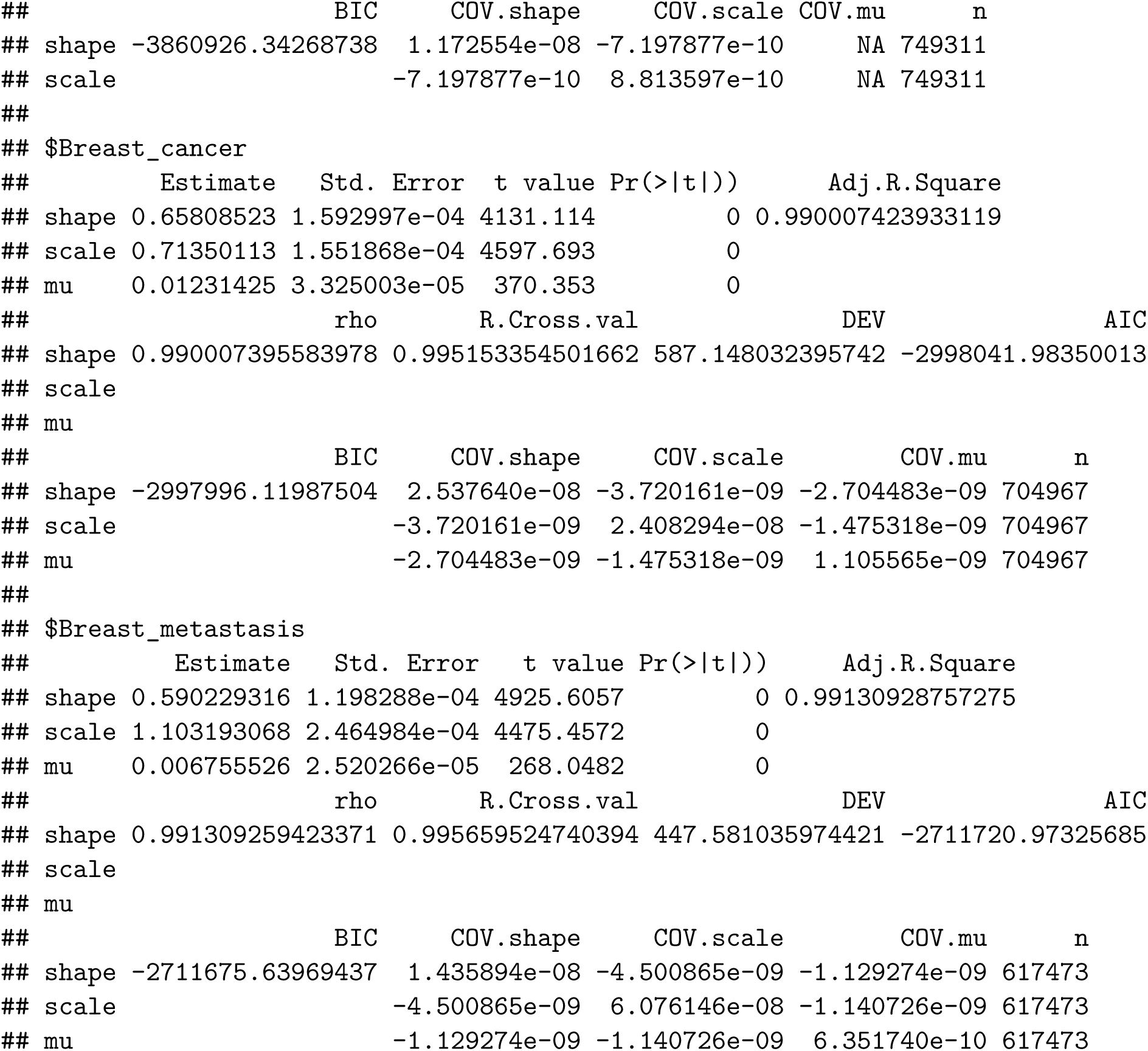

The goodness-of-fit indicators suggest that the fit to Weibull distribution model for *Ref0* is better than for *Ref*.

## 6 Signal detection

The information thermodynamics-based approach is postulated to provide greater sensitivity for resolving true signal from the thermodynamic background within the methylome (Sanchez and Mackenzie 2016). Because the biological signal created within the dynamic methylome environment characteristic of plants is not free from background noise, the approach, designated Methyl-IT, includes the application of signal detection theory (Greiner, Pfeiffer, and Smith 2000; Carter et al. 2016; Harpaz et al. 2013; Kruspe et al. 2017). Signal detection is a critical step to increase sensitivity and resolution of methylation signal by reducing the signal-to-noise ratio and objectively controlling the false positive rate and prediction accuracy/risk.

### 6.1 Potential methylation signal

The first estimation in our signal detection step is the identification of the cytosine sites carrying potential methylation signal *P S*. The methylation regulatory signal does not hold Weibull distribution and, consequently, for a given level of significance α (Type I error probability, e.g. *α* = 0.05), cytosine positions *k* with information divergence *H_k_ >*= *Hα*_=0.05_ can be selected as sites carrying potential signals *P S*. The value of α can be specified. For example, potential signals with *H_k_ > Hα*_=0.01_ can be selected. For each sample, cytosine sites are selected based on the corresponding fitted Weibull distribution model estimated in the previous step. Additionally, since cytosine with *T V*_*k*_ < 0.1 are the most abundant sites, depending on the sample (experiment), cytosine positions *k* with *H*_*k*_ > = *Hα*_=0.05_ and *T V*_*k*_ < 0.1 can be observed. To prevent the last situation we can select the *P S* with the additional constraint *T V*_*k*_ > T V_0_, where *T V*_0_ (‘tv.cut’) is a user specified value. The *P S* is detected with the function ‘getPotentialDIMP’:

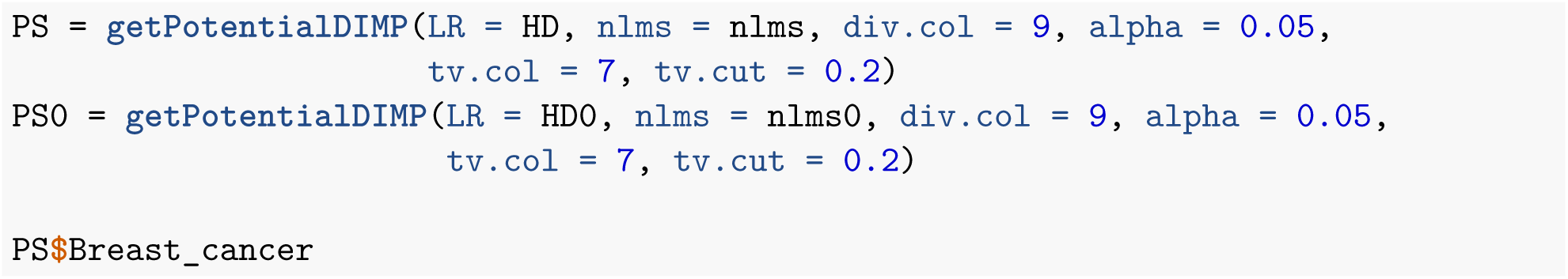

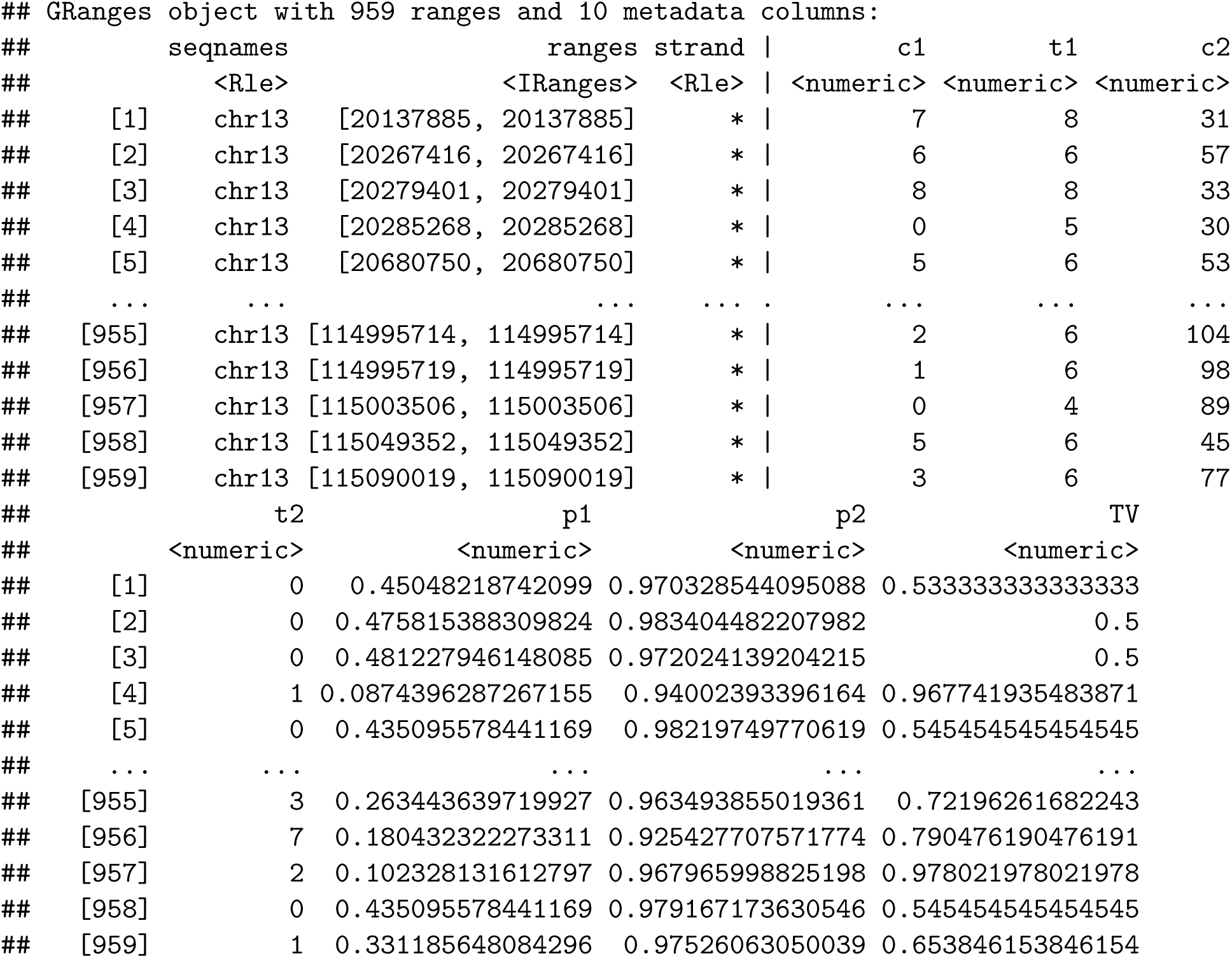
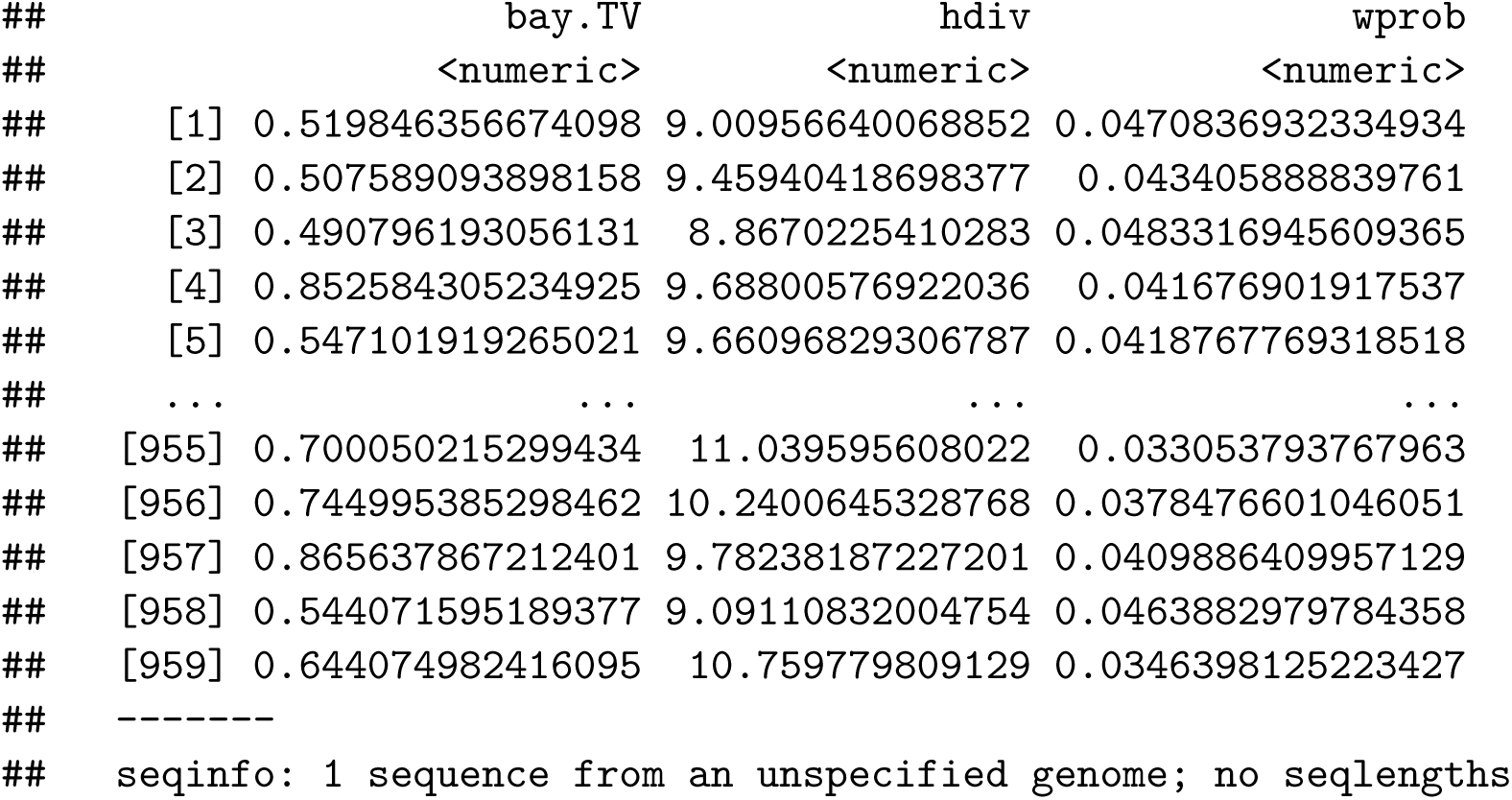

Notice that the total variation distance |*T V |* is an information divergence as well and it can be used in place of Hellinger divergence (Sanchez and Mackenzie 2016). The set of vectors *P*_*i*_ = (*p*_*i*_, 1 − *p*_*i*_) and distance function |*T V |* integrate a metric space. In particular:

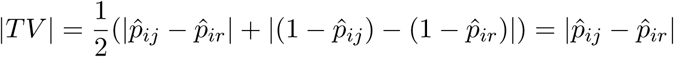

That is, the quantitative effect of the vector components 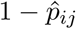 and 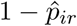 (in our case, the effect of unmethylated read counts) is not present in *T V* as in 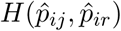.

### 6.2 Histogram and boxplots of methylation potential signals

As before, a single GRanges object is built from the above set GRanges objects using the function ‘uniqueGRanges’, and the Hellinger divergences of the cytosine sites carrying *P S* (for each sample) are located in a single matrix on the metacolumn of the GRanges object.

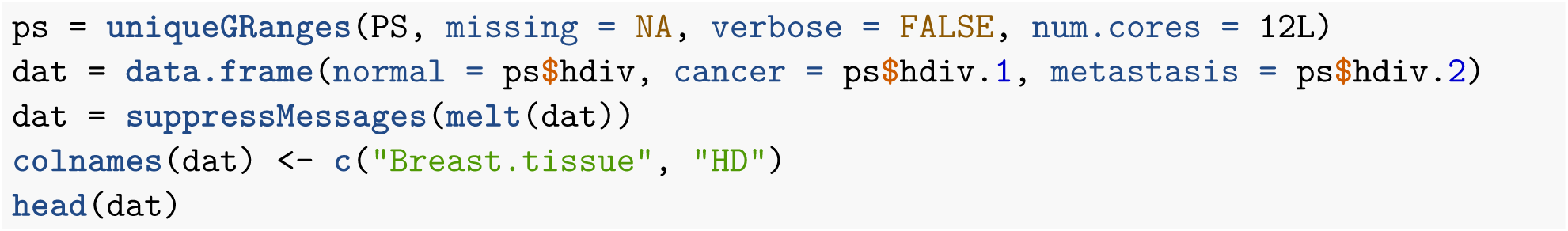

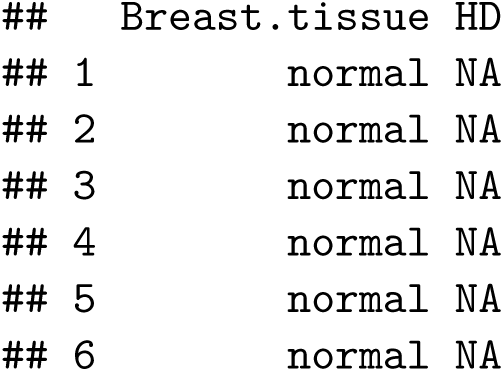

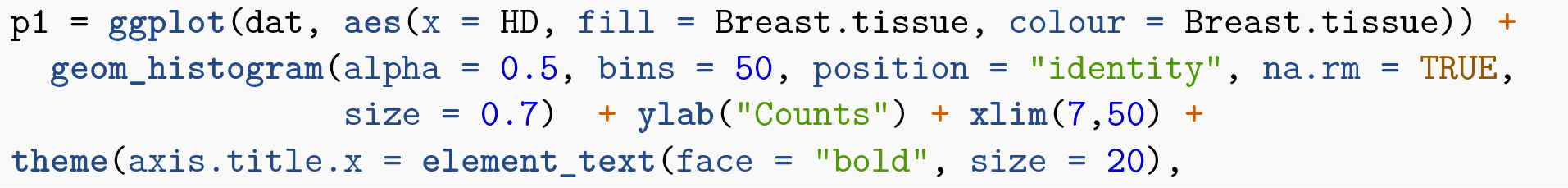
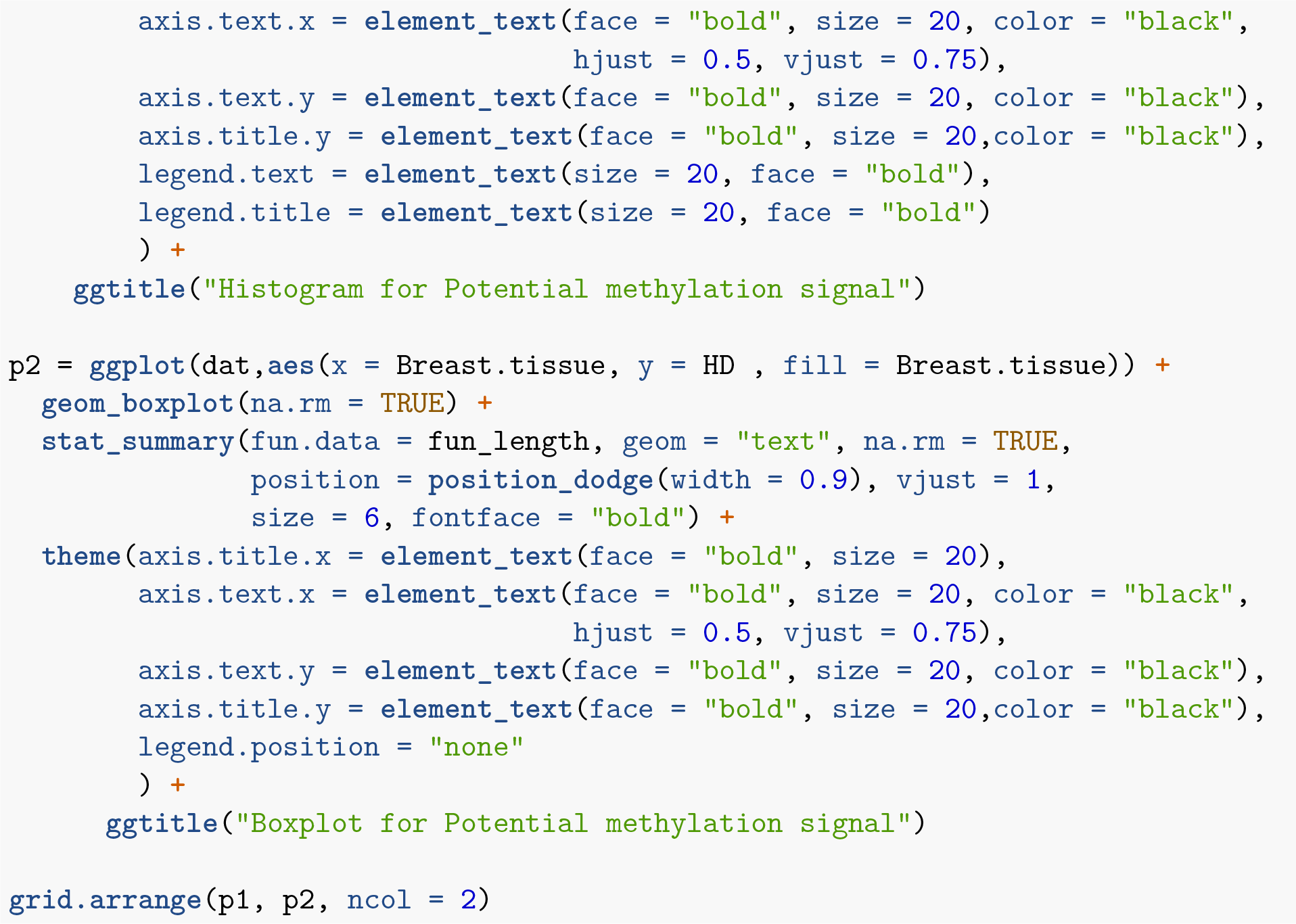

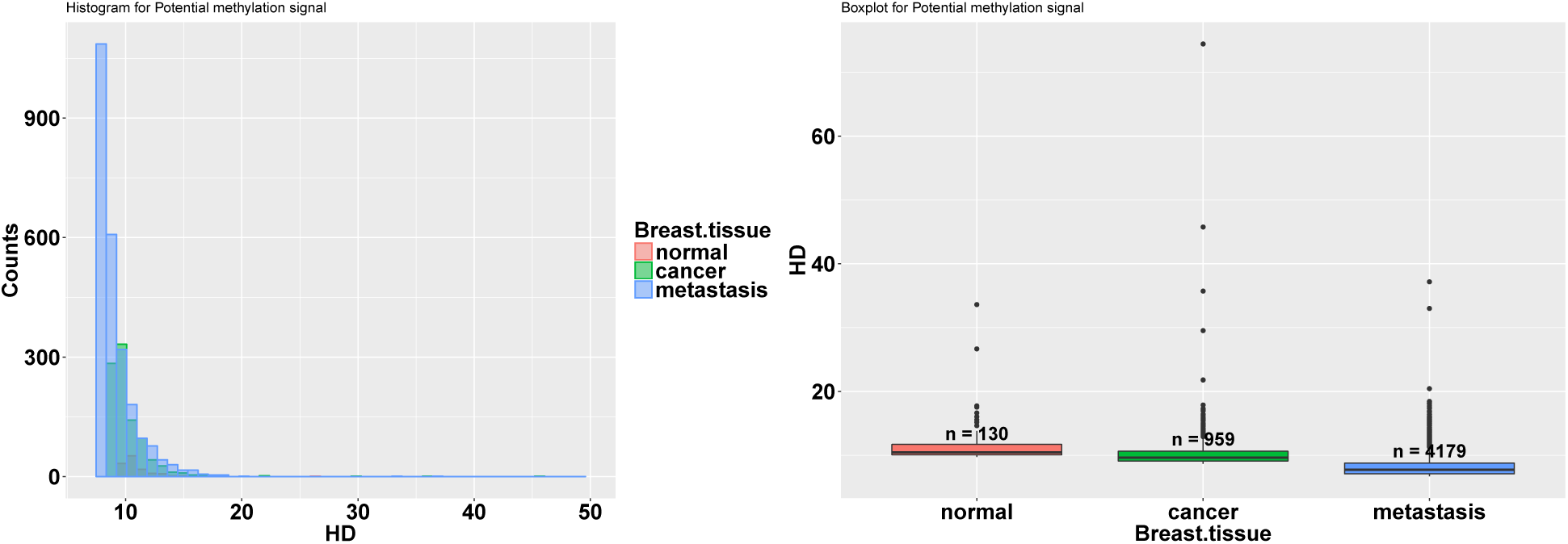

## 7 Cutpoint estimation

Laws of statistical physics can account for background methylation, a response to thermal fluctuations that presumably functions in DNA stability (Sanchez and Mackenzie 2016). True signal is detected based on the optimal cutpoint (López-Ratón et al. 2014), which can be estimated from the area under the curve (AUC) of a receiver operating characteristic (ROC) curve built from a logistic regression performed with the potential signals from controls and treatments. The ROC AUC is equivalent to the probability that a randomly-chosen positive instance is ranked more highly than a randomly-chosen negative instance (Fawcett 2005). In the current context, the AUC is equivalent to the probability to distinguish a randomly-chosen methylation regulatory signal induced by the treatment from a randomly-chosen signal in the control.

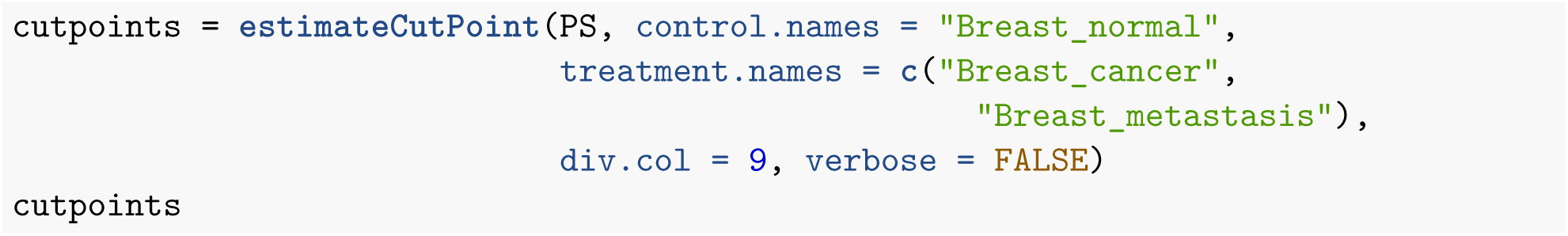

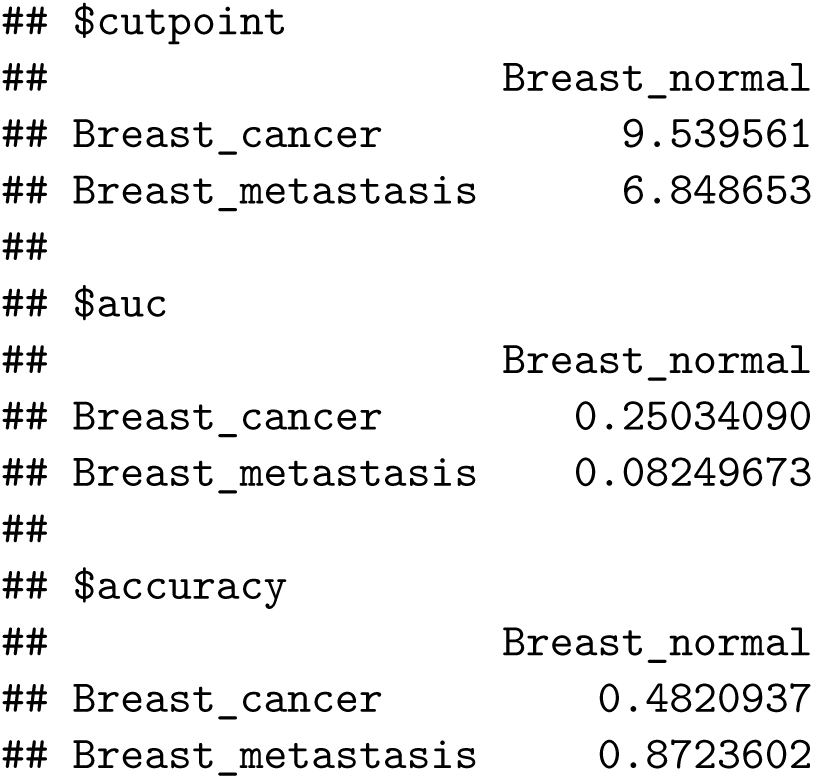

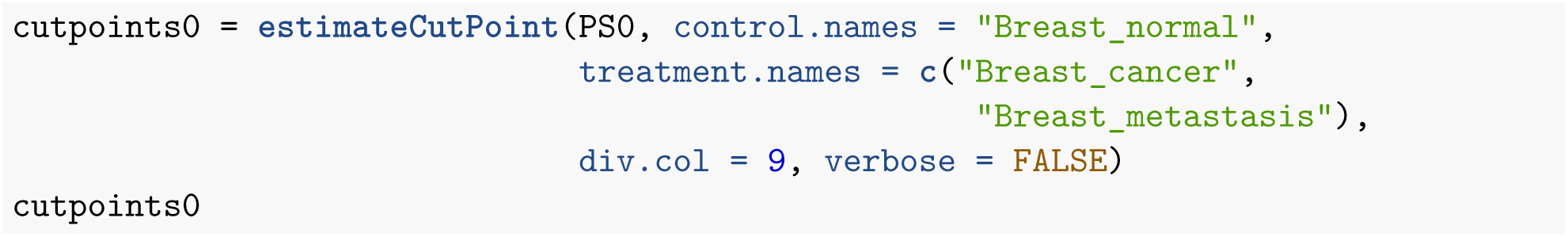

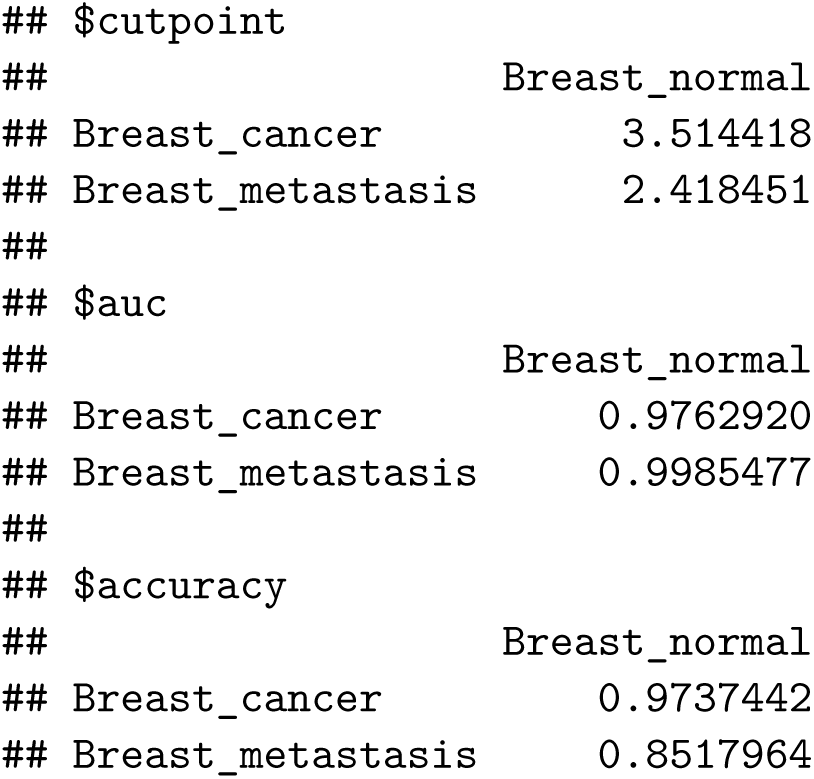

In practice, potential signals are classified as “control”’ (*CT*) and “treatment”’ (*T T*) signals (prior classification) and the logistic regression (LG): signal (with levels *CT* (0) and *T T* (1)) versus *H_k_* is performed. LG output yields a posterior classification for the signal. Prior and posterior classifications are used to build the ROC curve and then to estimate AUC and cutpoint *H_cutpoint_*.

## 8 DIMPs

Cytosine sites carrying a methylation signal are designated *differentially informative methylated positions* (DIMPs). The probability that a DIMP is not induced by the treatment is given by the probability of false alarm (*P_F A_*, false positive). That is, the biological signal is naturally present in the control as well as in the treatment. Each DIMP is a cytosine position carrying a significant methylation signal, which may or may not be represented within a differentially methylated position (DMP) according to Fisher’s exact test (or other current tests). A DIMP is a DNA cytosine position with high probability to be differentially methylated or unmethylated in the treatment with respect to a given control. Notice that the definition of DIMP is not deterministic in an ordinary sense, but stochastic-deterministic in physico-mathematical terms.

DIMPs are selected with the function:

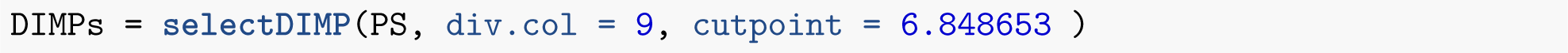

### 8.1 Histogram and boxplots of DIMPs

The cutpoint detected with the signal detection step is very close (in this case) to the Hellinger divergence value *H*_*α=0.05*_ estimated for cancer tissue. The natural methylation regulatory signal is still present in a patient with cancer and reduced during the metastasis step. This signal is detected here as a false alarm (*P*_*F A*_, false positive)

The list of GRanges with DIMPs are integrated into a single GRanges object with the matrix of ‘hdiv’ values on its metacolumn:

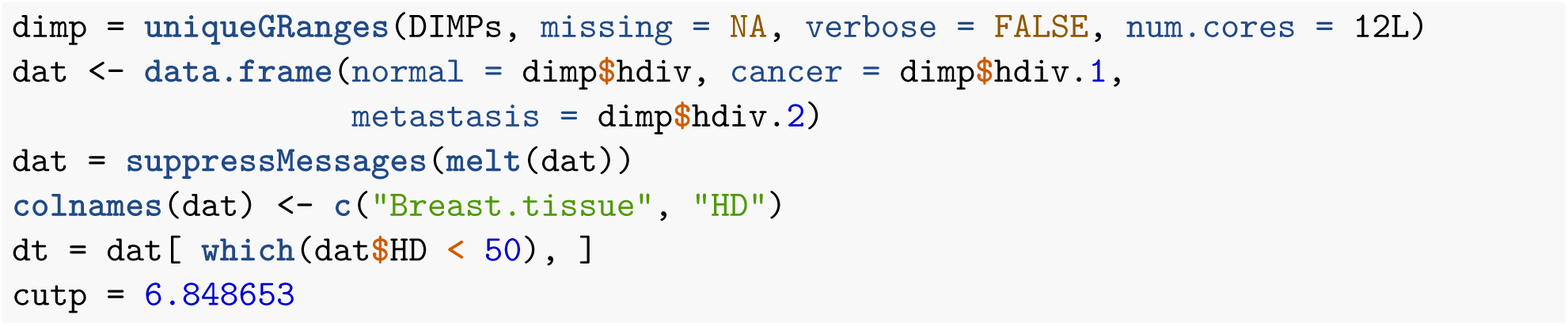

The multiplot with the histogram and the boxplot can now built:

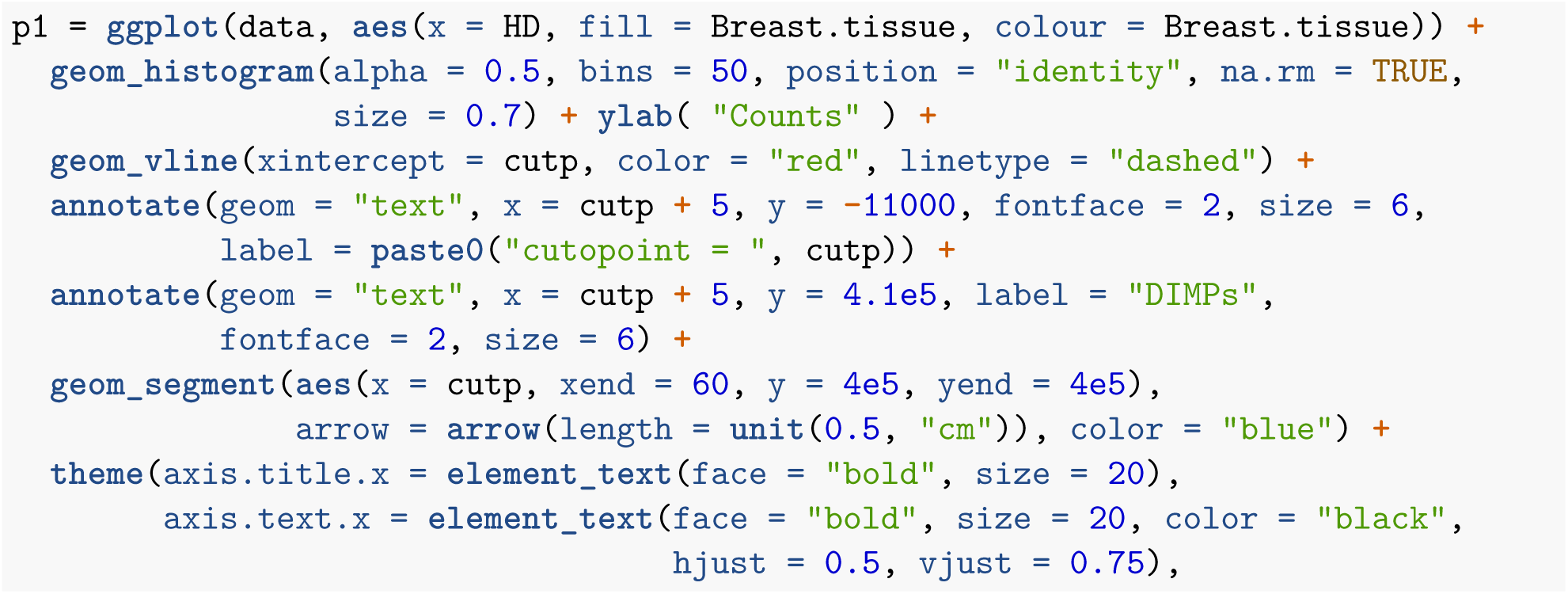
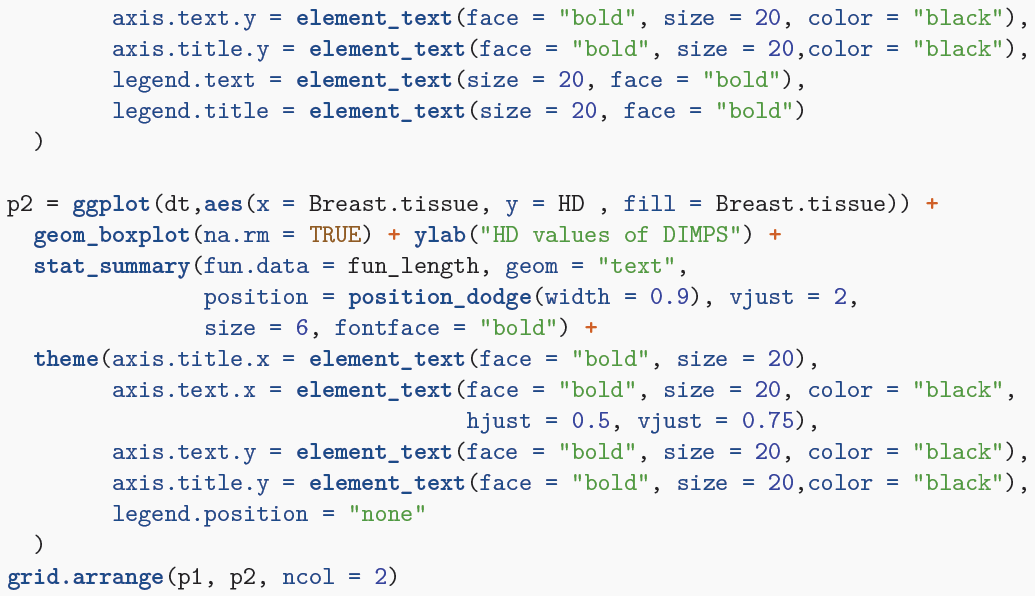

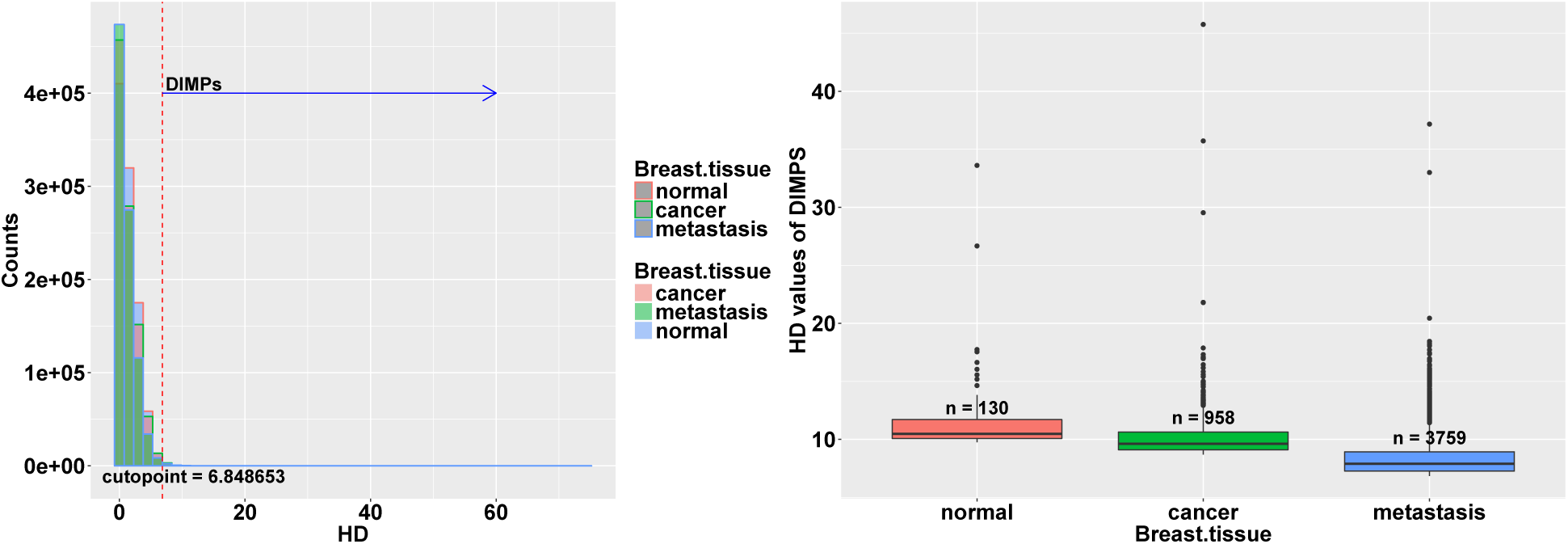

### 8.2 Venn Diagram of DIMPs

The Venn diagram of DIMPs reveals that the number cytosine site carrying methylation signal with a divergence level comparable to that observed in breast tissues with cancer and metastasis is relatively small (2797 DIMPs). The number of DIMPs decreased in the breast tissue with metastasis, but, as shown in the last boxplot, the intensity of the signal increased.

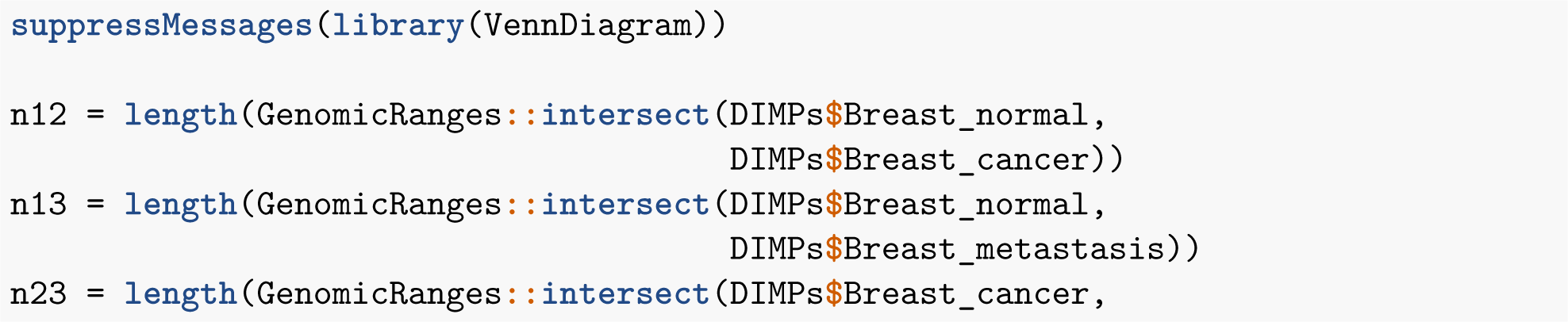
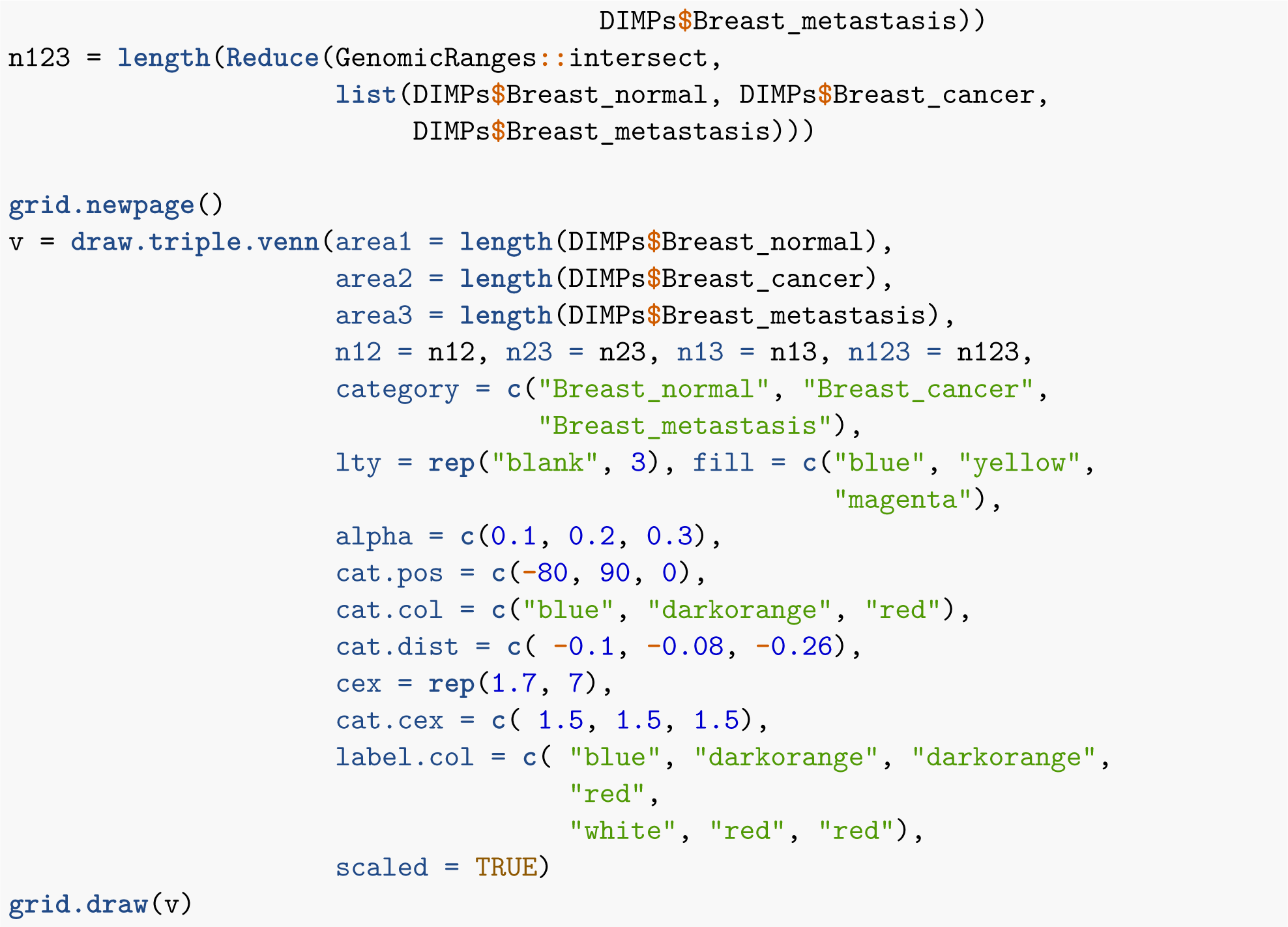

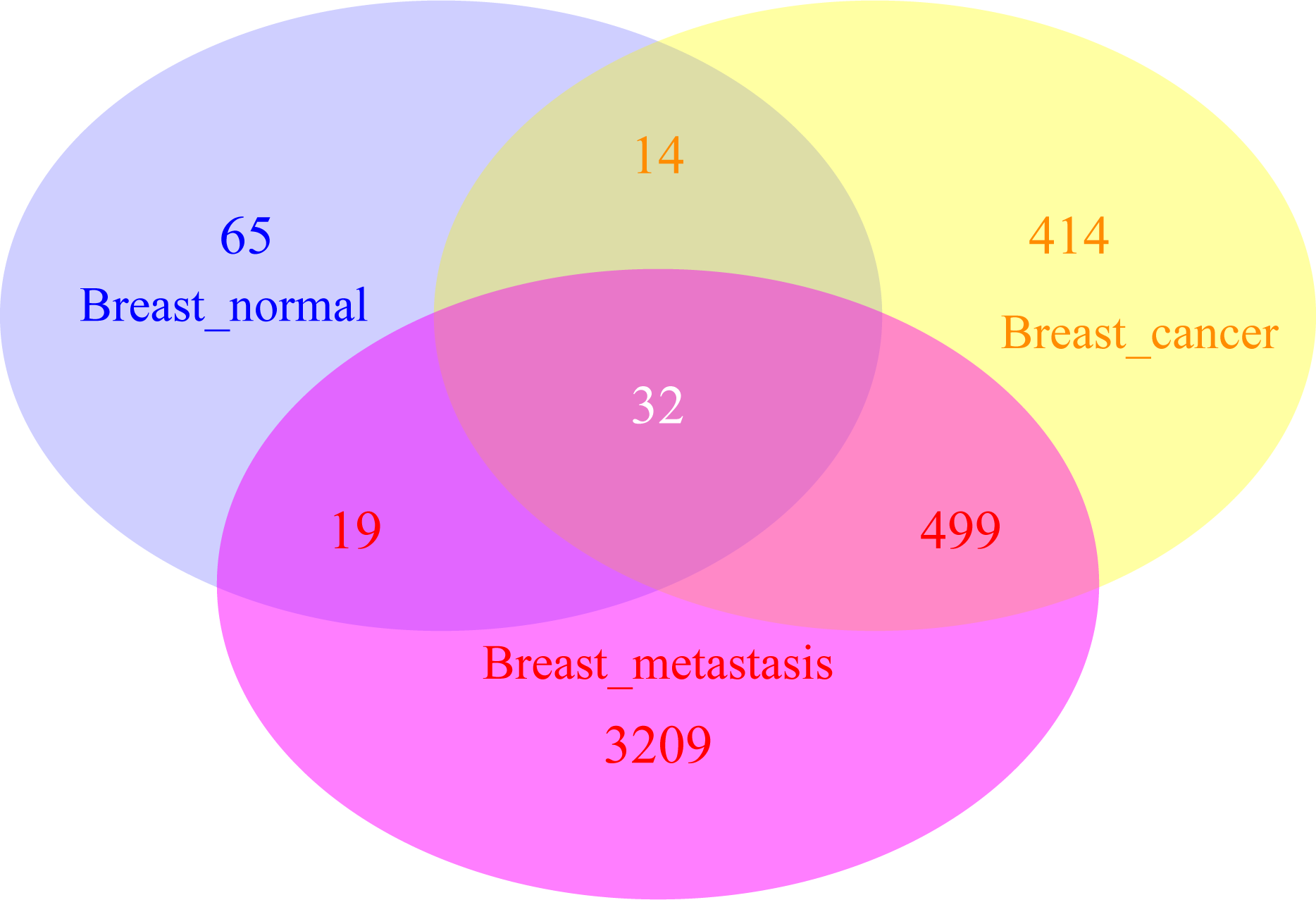

Notice that natural methylation regulatory signals (not induced by the treatment) are present in both groups, control and treatment. The signal detection step permits us to discriminate the “ordinary” signals observed in the control from those induced by the treatment (a disease in the current case). In addition, this diagram reflects a classification of DIMPs only based on the cytosine positions. That is, this Venn diagram cannot tell us whether DIMPs at the same position can be distinguishable or not. For example, DIMPs at the same positions in control and treatment can happened with different probabilities estimated from their corresponding fitted Weibull distributions (see below).

### 8.3 Venn Diagram of *DIMPs for reference Ref0*

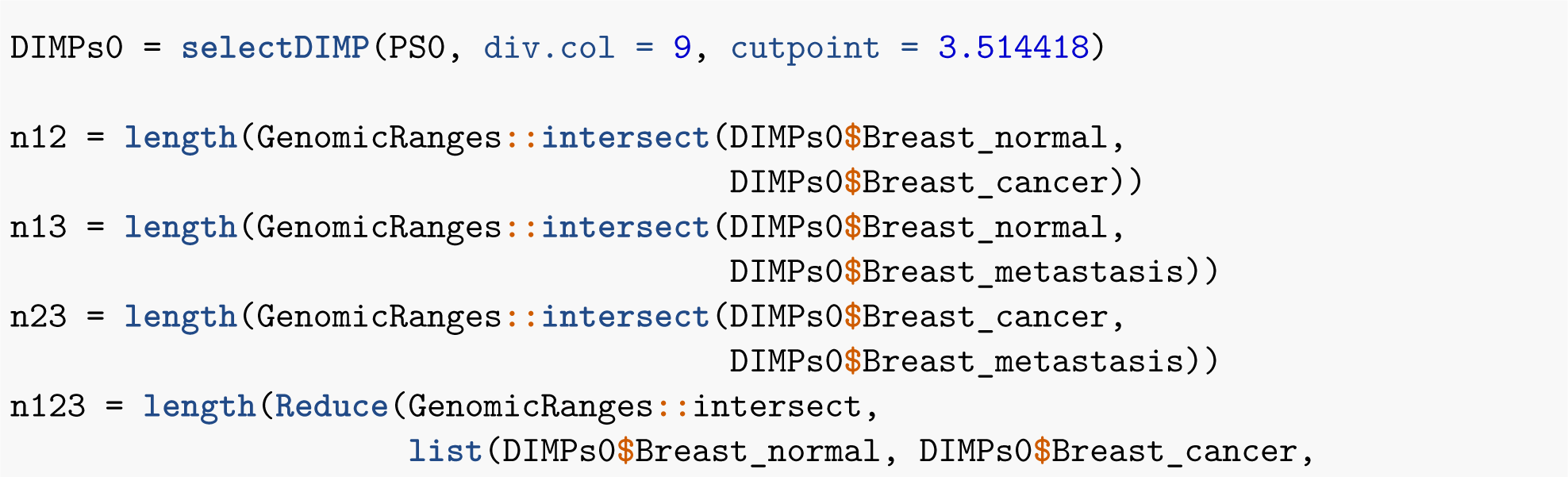

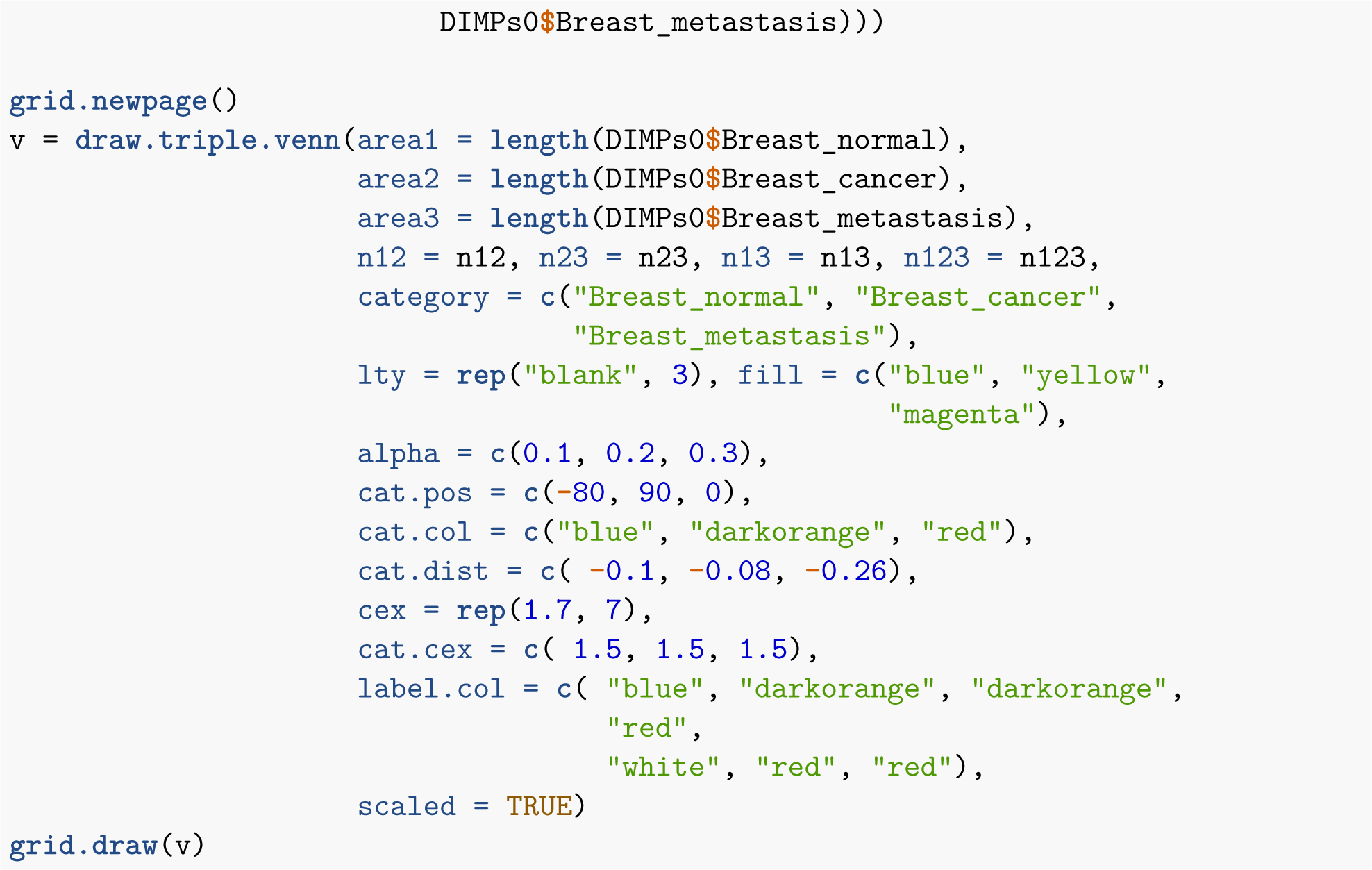

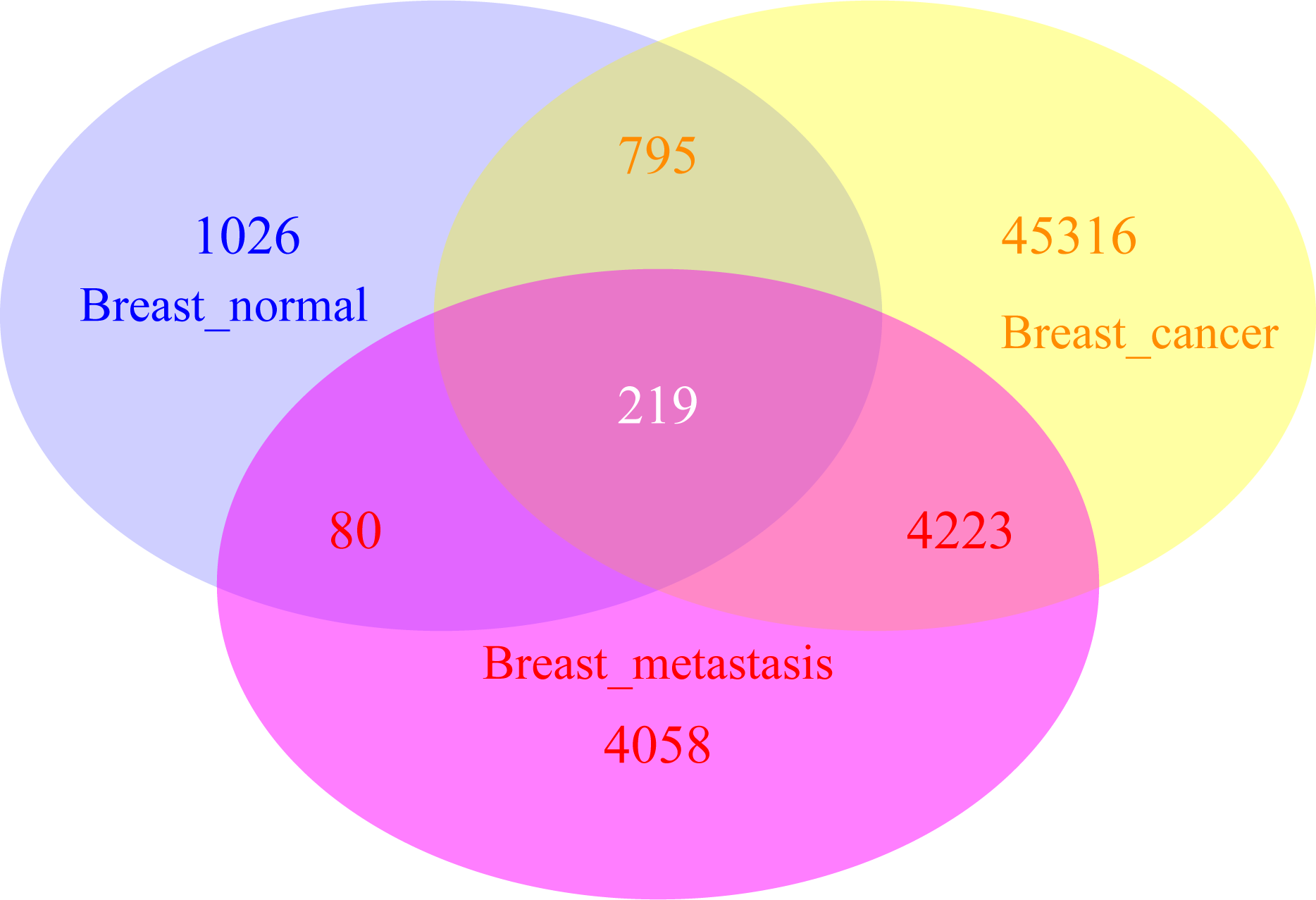

## 9 Differentially informative methylated genomic regions (DIMRs)

Our degree of confidence in whether DIMP counts in both groups of samples, control and treatment, represent true biological signal was determined in the signal detection step. To estimate DIMRs, we followed similar steps to those proposed in Bioconductor R package DESeq2 (Love, Huber, and Anders 2014), but our GLM test looks for statistical difference between the groups based on gene-body DIMP counts overlapping a given genomic region rather than read counts. The regression analysis of the generalized linear model (GLM) with logarithmic link was applied to test the difference between group counts. The fitting algorithmic approaches provided by ‘glm’ and ‘glm.nb’ functions from the R packages stat and MASS, respectively, were used for Poisson (PR), Quasi-Poisson (QPR) and Negative Binomial (NBR) linear regression analyses, respectively.

## 9.1 Differentially methylated genes (DMGs)

We shall call DMGs those DIMRs restricted to gene-body regions. DMGs are detected using function ‘countTest’. We used computational steps from DESeq2 packages. In the current case we follow the steps:

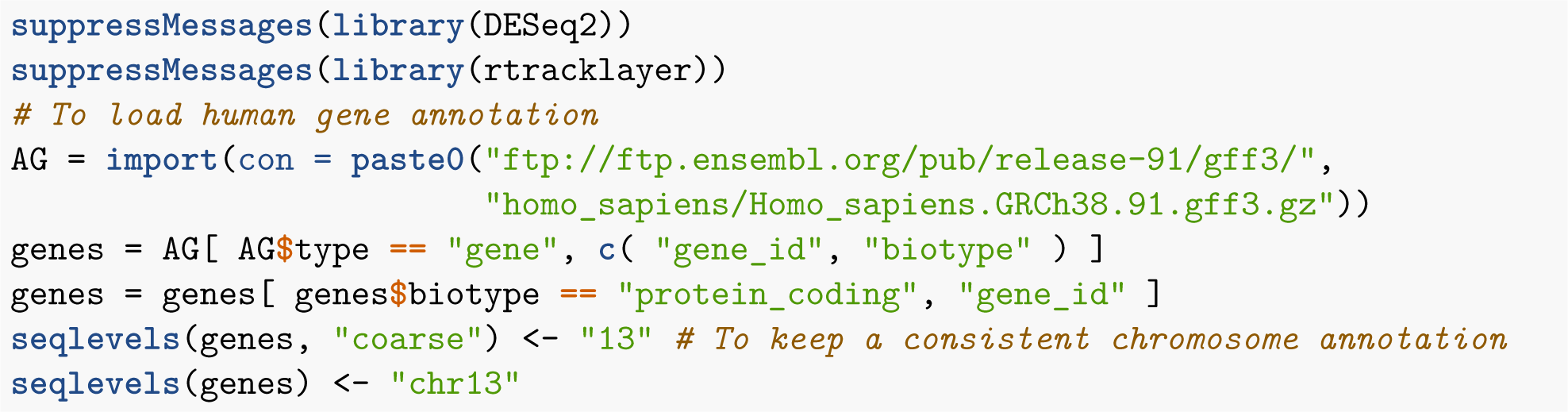

Function ‘getDIMPatGenes’ is used to count the number of DIMPs at gene-body. The operation of this function is based on the ‘findOverlaps’ function from the ‘GenomicRanges’ Bioconductor R package. The ‘findOverlaps’ function has several critical parameters like, for example, ‘maxgap’, ‘minoverlap’, and ‘ignore.strand’. In our function ‘dimpAtGenes’, except for setting ignore.strand = TRUE and type = “within”, we preserve the rest of default ‘findOverlaps’ parameters. In this case, these are important parameter settings because the local mechanical effect of methylation changes on a DNA region where a gene is located is independent of the strand where the gene is encoded. That is, methylation changes located in any of the two DNA strands inside the gene-body region will affect the flexibility of the DNA molecule (Choy et al. 2010; Severin et al. 2011).

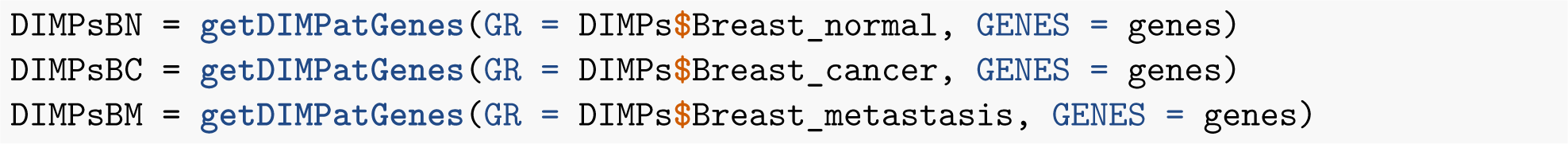

The number of DIMPs on the strand where a gene is encoded is obtained by setting ignore.strand = FALSE. However, for the current example results will be the same since the datasets downloaded from GEO do not have strand information. Next, the above GRanges objects carrying the DIMP counts from each sample are grouped into a single GRanges object. Since we have only one control, to perform group comparison and to move forward with this example, we duplicated ‘Breast_normal’ sample. Obviously, the confidence on the results increases with the number of sample replications per group (in this case, it is only an illustrative example on how to perform the analysis, since a fair comparison requires for more than one replicate in the control group).

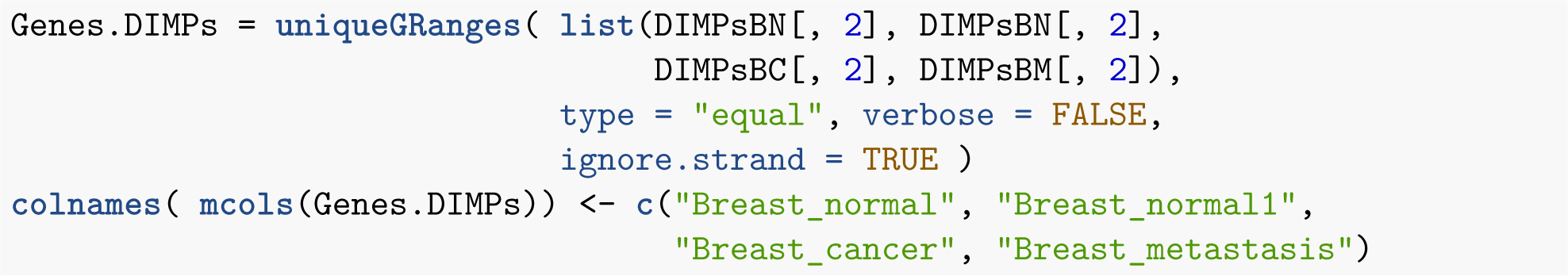

Next, the set of mapped genes are annotated

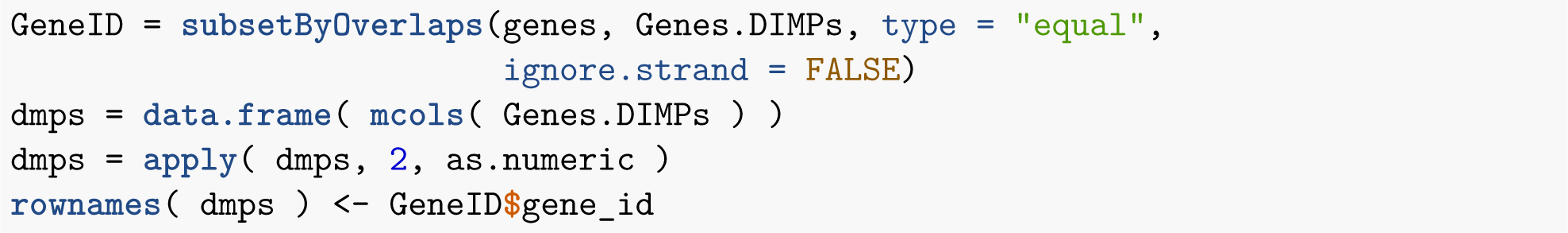

Now, we build a ‘DESeqDataSet’ object using functions DESeq2 package.

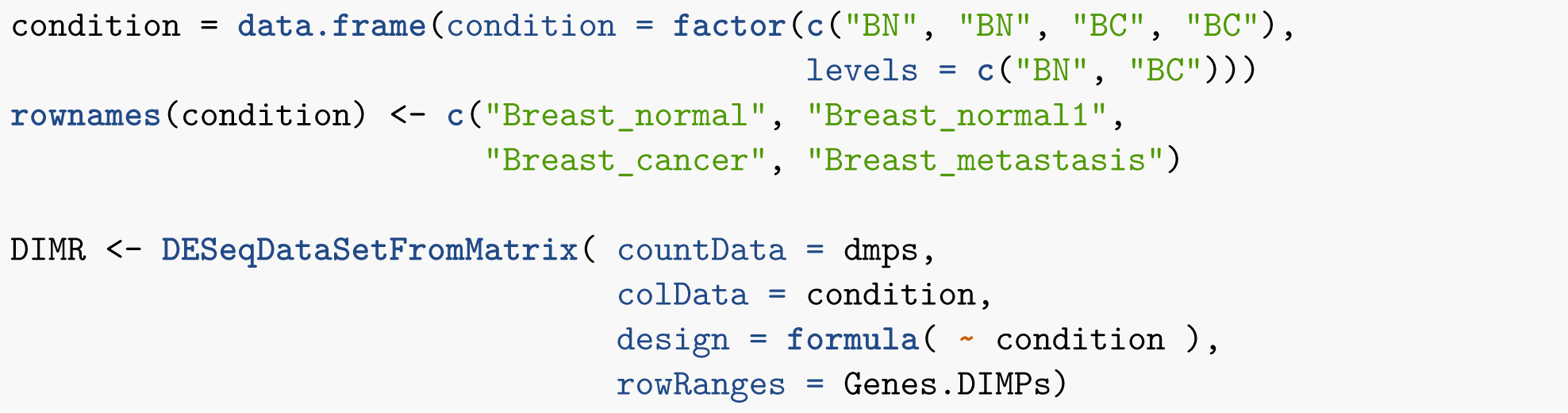

`## converting counts to integer mode`

DMG analysis is performed with the function ’countTest’

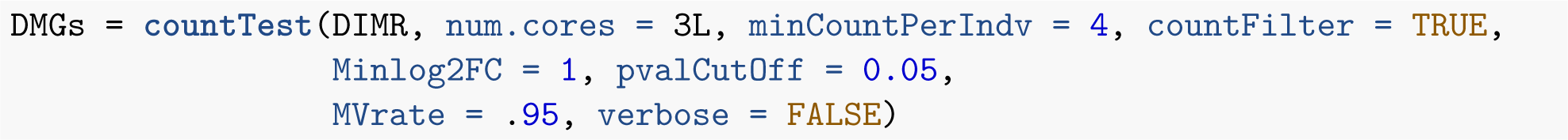

`## gene-wise dispersion estimates`

`## mean-dispersion relationship`

`## final dispersion estimates`

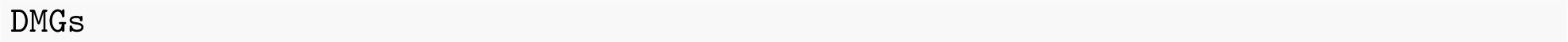

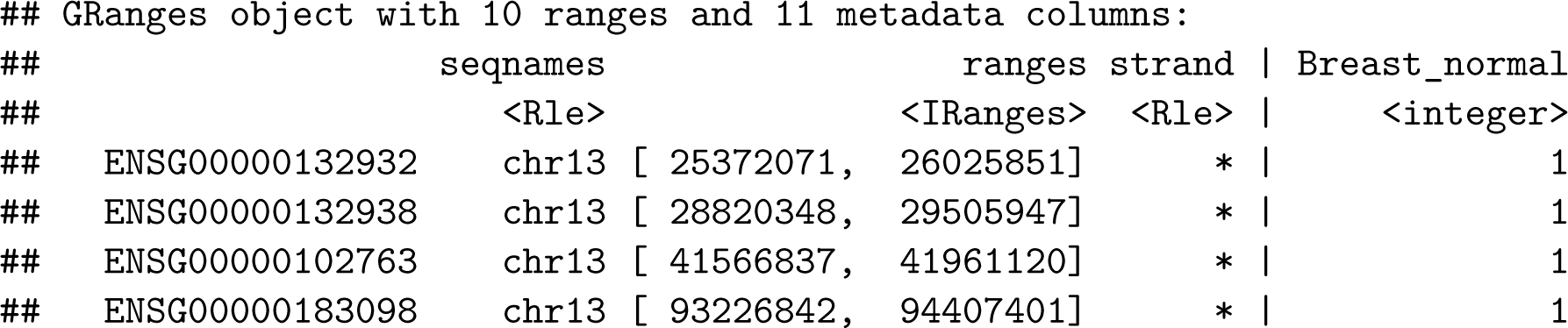
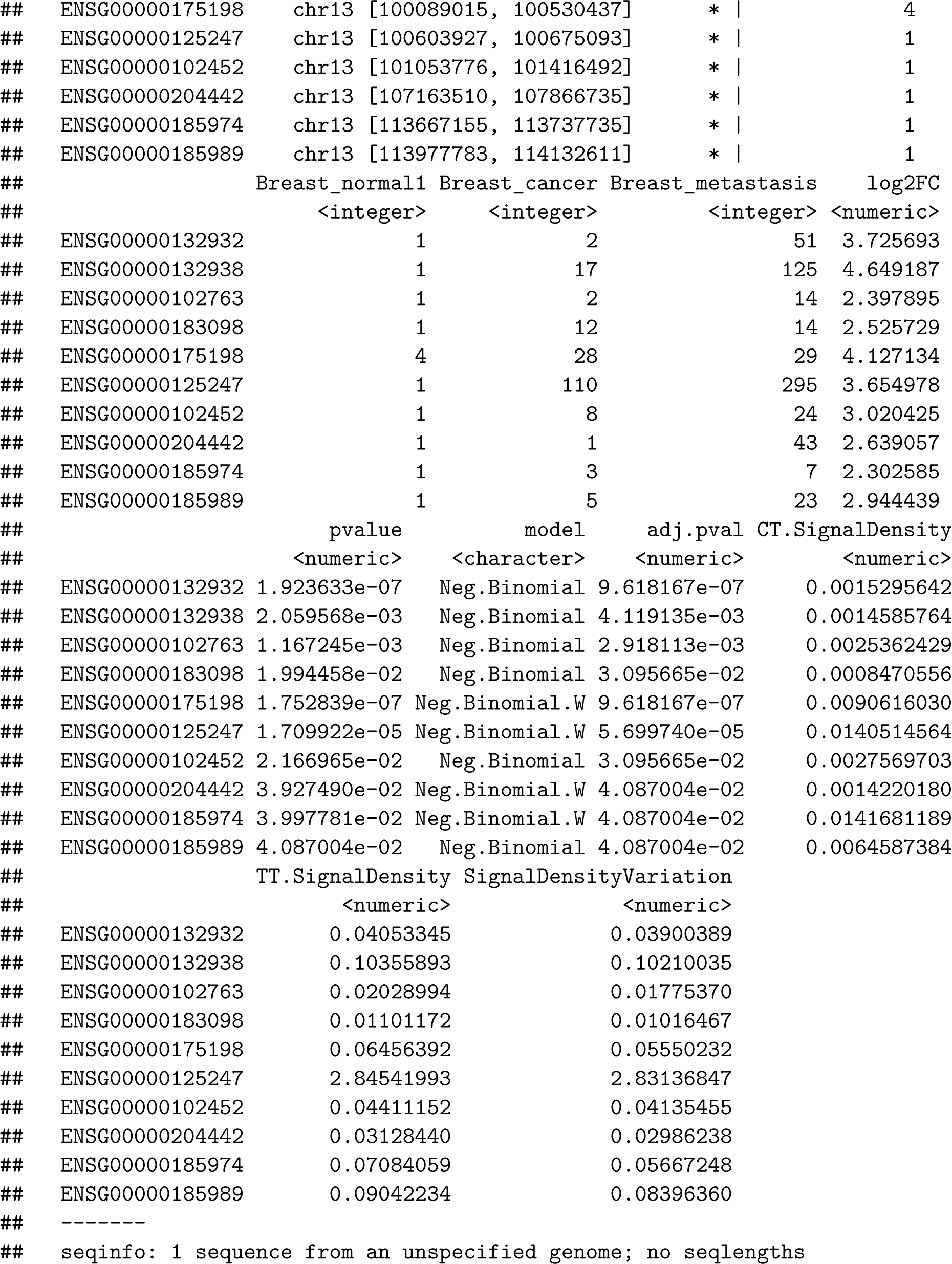

## 9.2. DMGs for reference Ref0

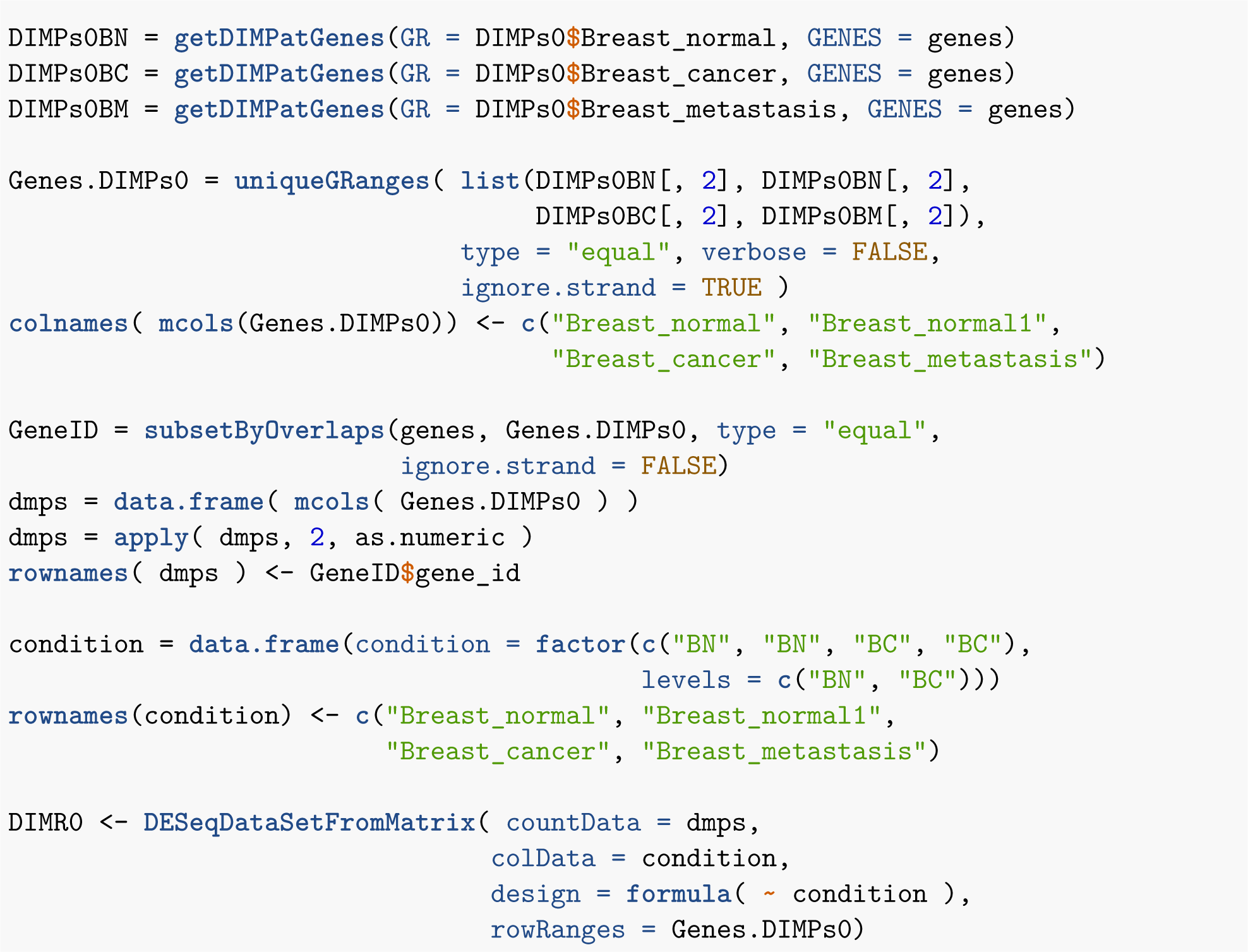

`## converting counts to integer mode`

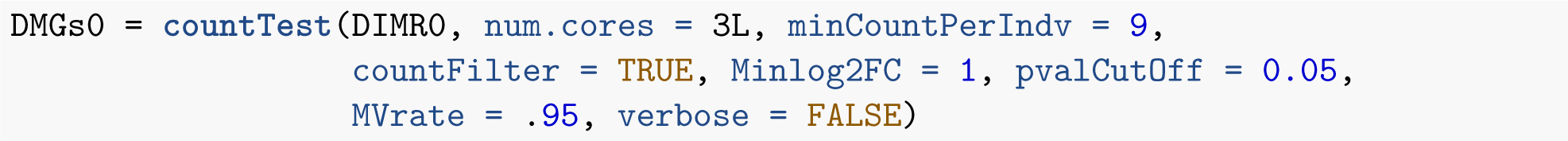

`## gene-wise dispersion estimates`

`## mean-dispersion relationship`

`## final dispersion estimates`

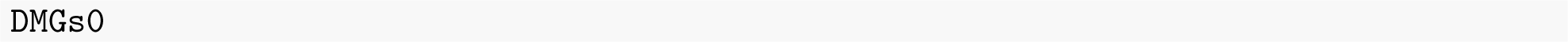

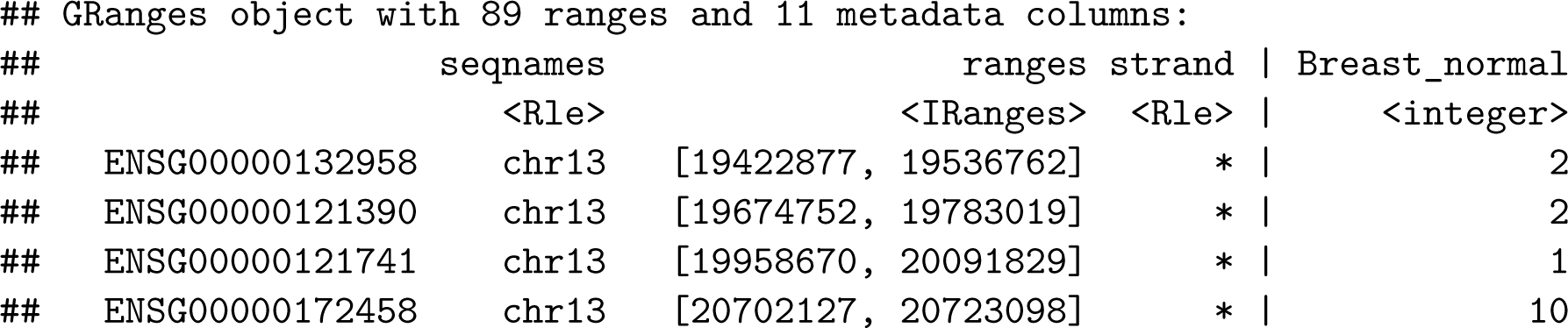
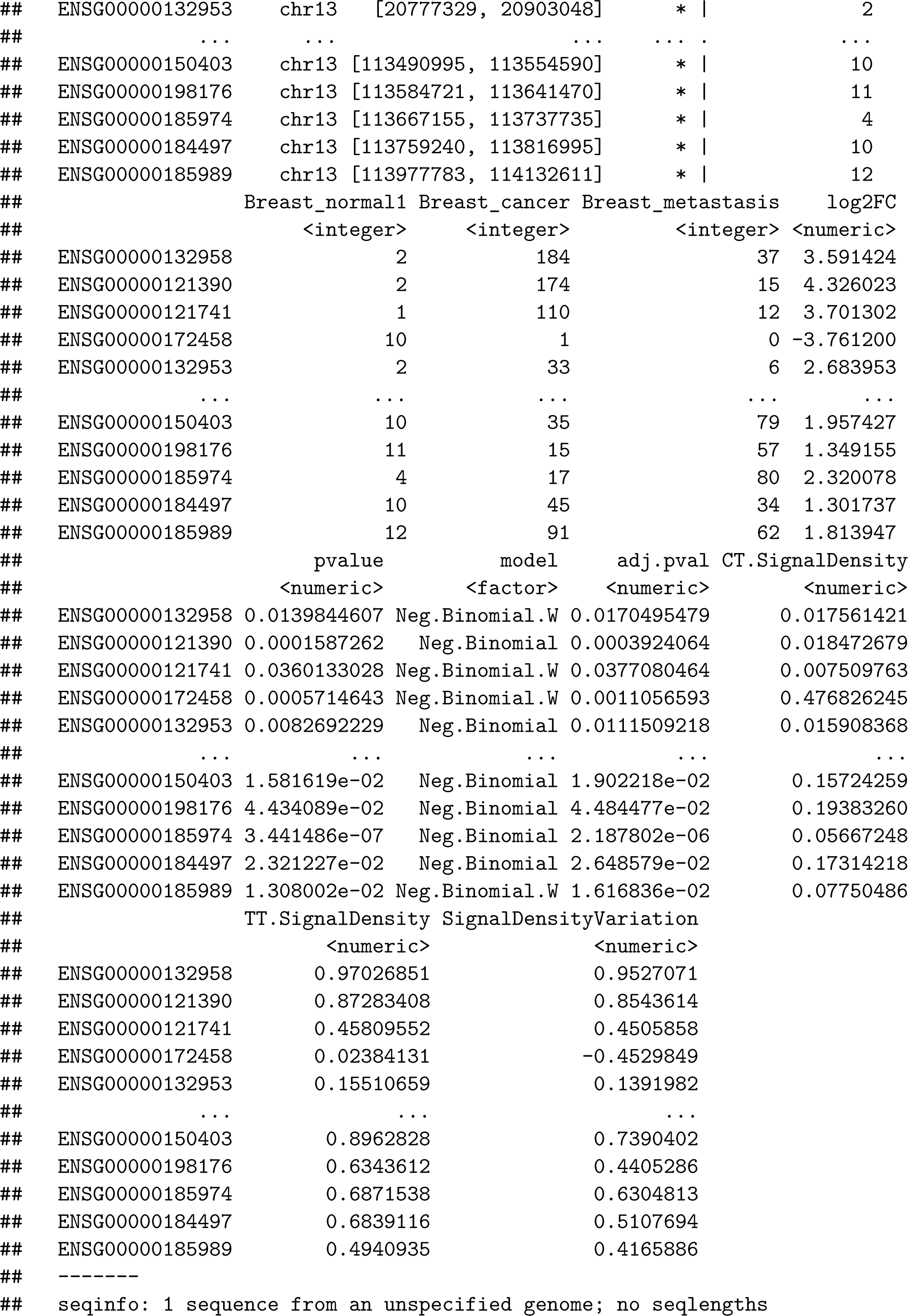

*BRCA2, a breast cancer associated risk gene, is found between the DMGs*

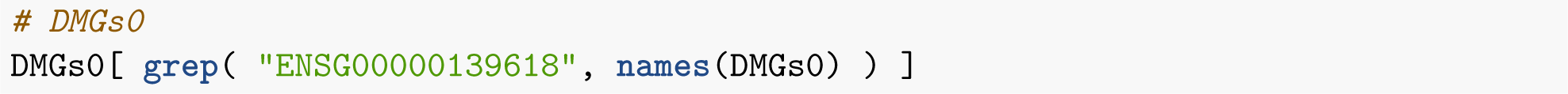

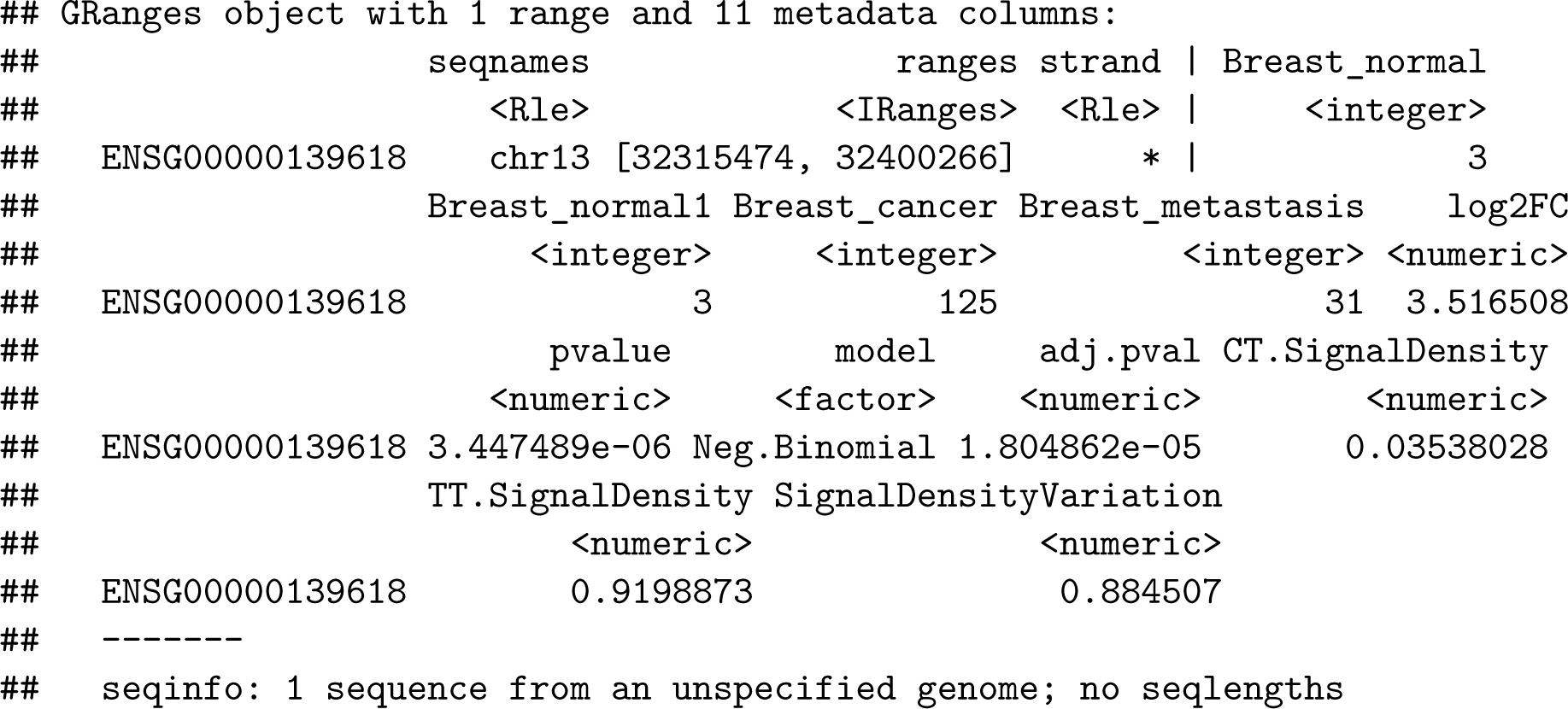

## 10 Classification of DIMPs into two classes

The regulatory methylation signal is an output from a natural process that continuously takes place across the ontogenetic development of the organism. Therefore, we expect to see methylation signal in natural, ordinary conditions. Function ‘evaluateDIMPclass’ can be used to perform a classification of DIMPs into two classes: DIMPS from control and DIMPs from treatment samples, as well as an evaluation of the classification performance (for more details see ?evaluateDIMPclass). In the setting below, a logistic regression: group versus divergence (at DIMPs), will be executed after randomly splitting the original DIMP dataset into two subsets: training (60%) and testing (40%).

The performance of the logistic classifier using reference ‘Ref’ is:

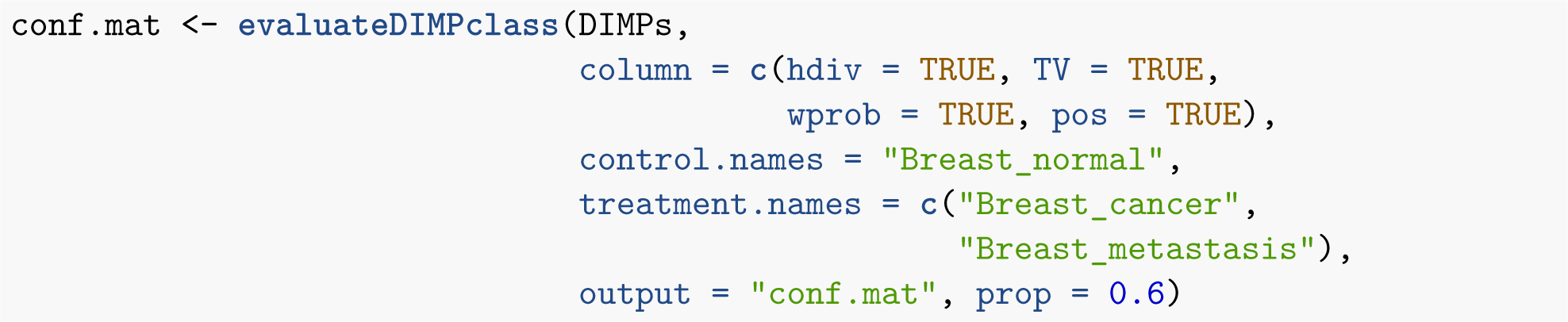

`## Model: treat ~ hdiv + TV + logP + pos`

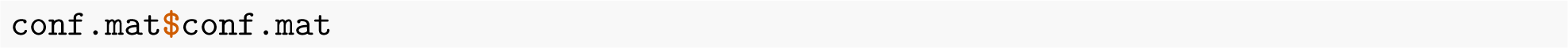

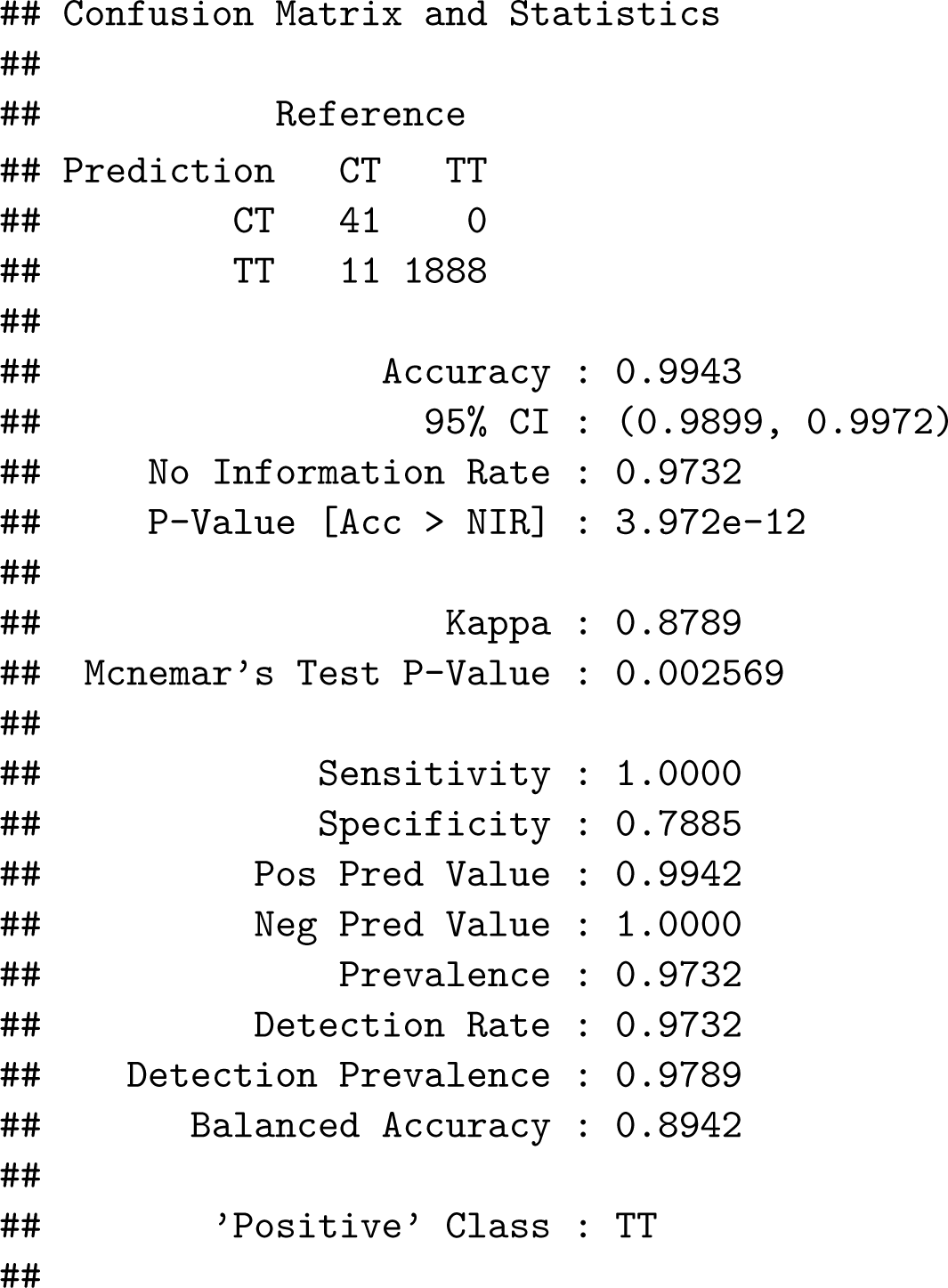

The best fitted logistic model using reference ‘Ref’ is:

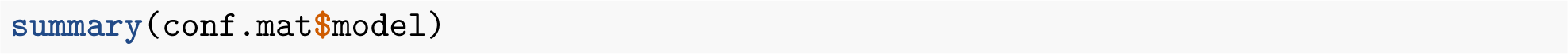

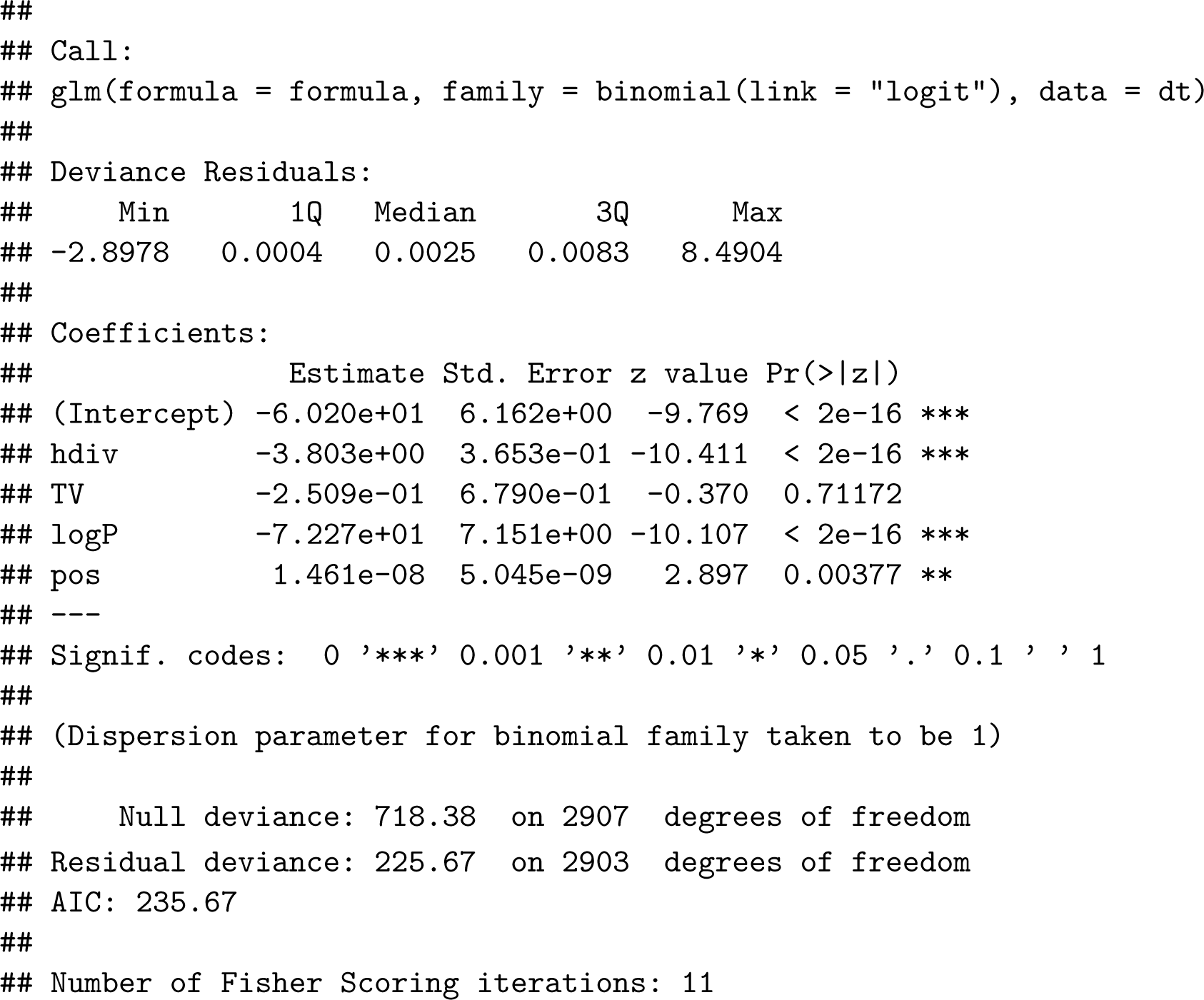

In this case, the only variable not included in the model is total variation *T V* and all the rest are significant. The generalized linear regression can be performed by removing the variables *T V*. There are three other classifiers available: “pca.logistic”, “pca.lda”, and “pca.qda” (type ?evaluateDIMPclass in R console for more details). Principal component analysis (PCA) is used to convert a set of observations of possibly correlated predictor variables into a set of values of linearly uncorrelated variables (principal components, PCs). Then, the PCs are used as new uncorrelated predictor variables for LDA, QDA, and logistic classifiers. In the current case, the best classification result is obtained with the combination PCA + Quadratic Discriminant Analysis (PCA + QDA, “pca.qda”).

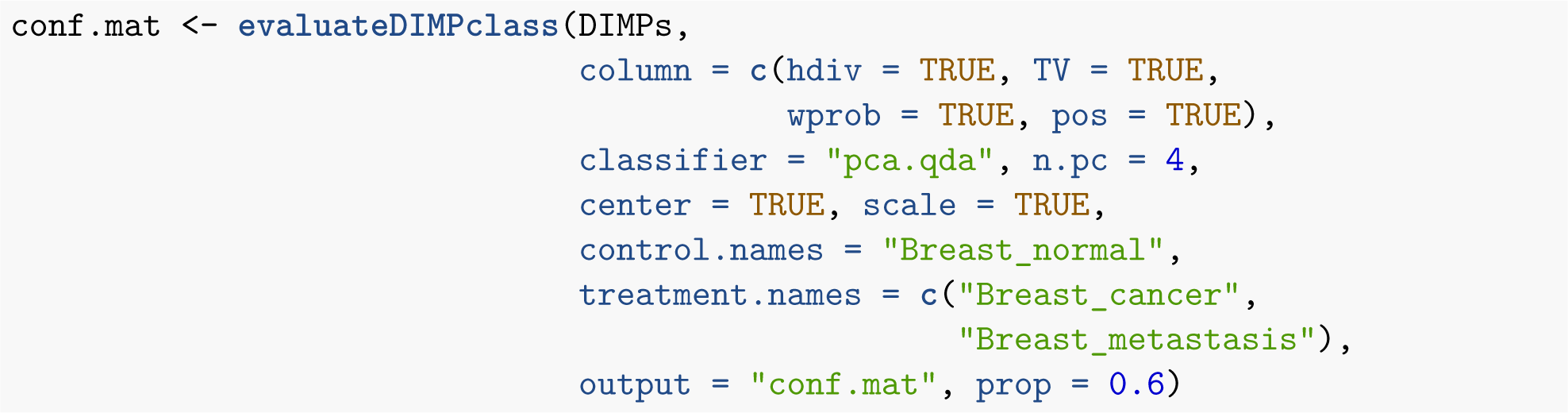

`## Model: treat ~ hdiv + TV + logP + pos`

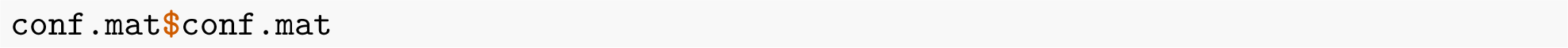

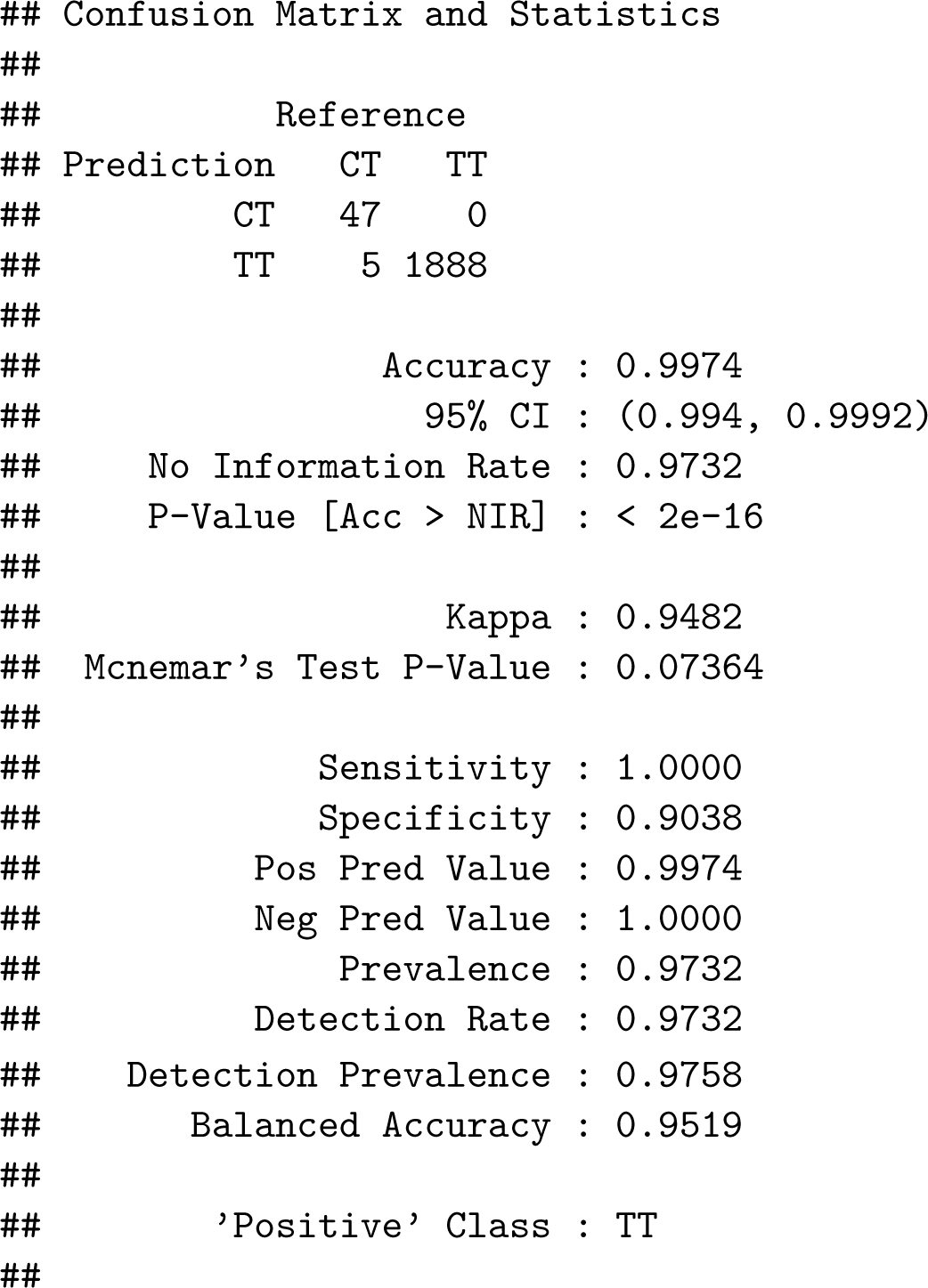

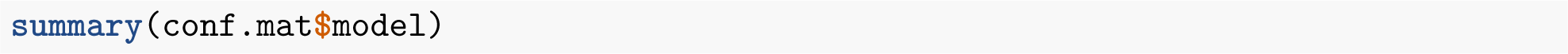

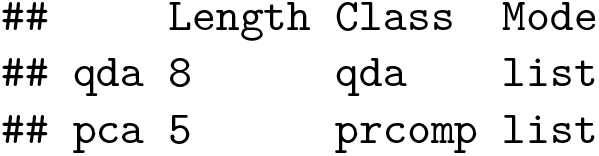

Monte Carlo (bootstrap) validation with 500 resamplings is performed by using the option ‘output = “mc.val”’:

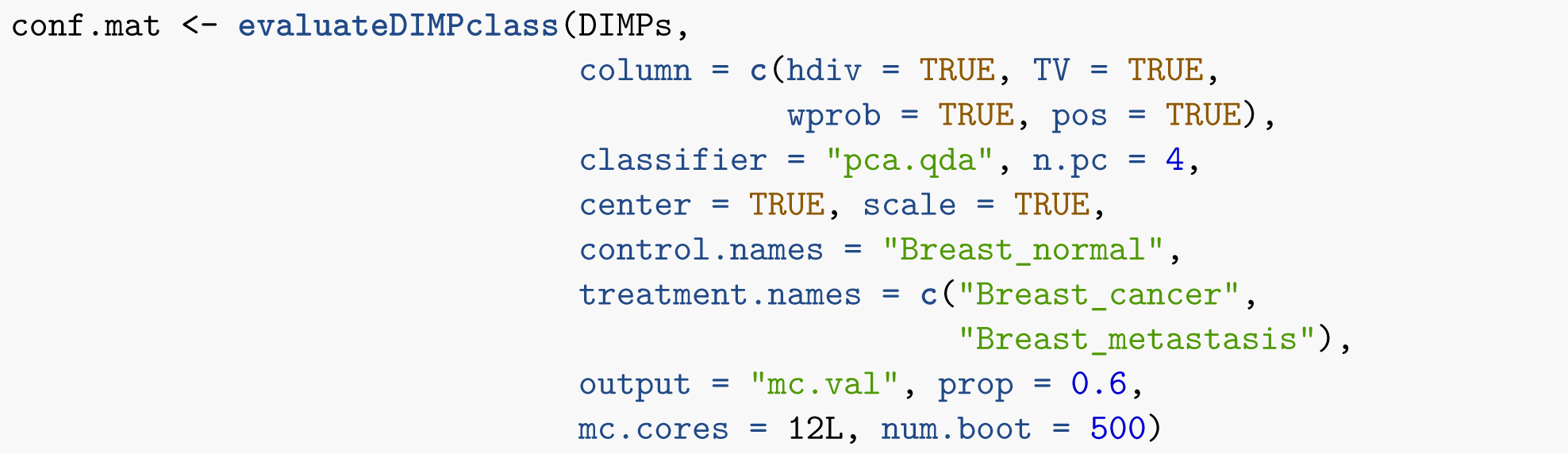

`## Model: treat ~ hdiv + TV + logP + pos`

`conf.mat`

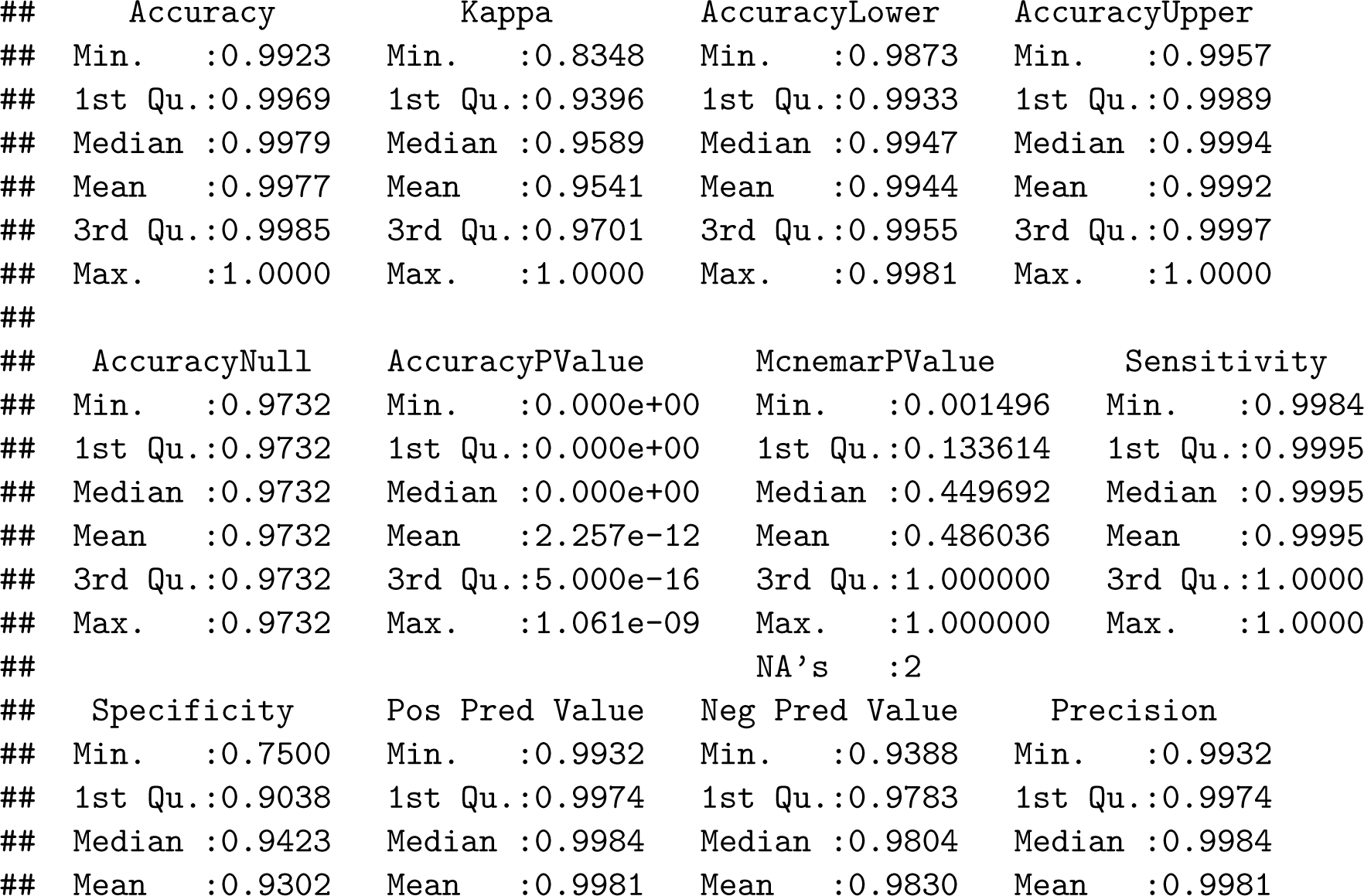
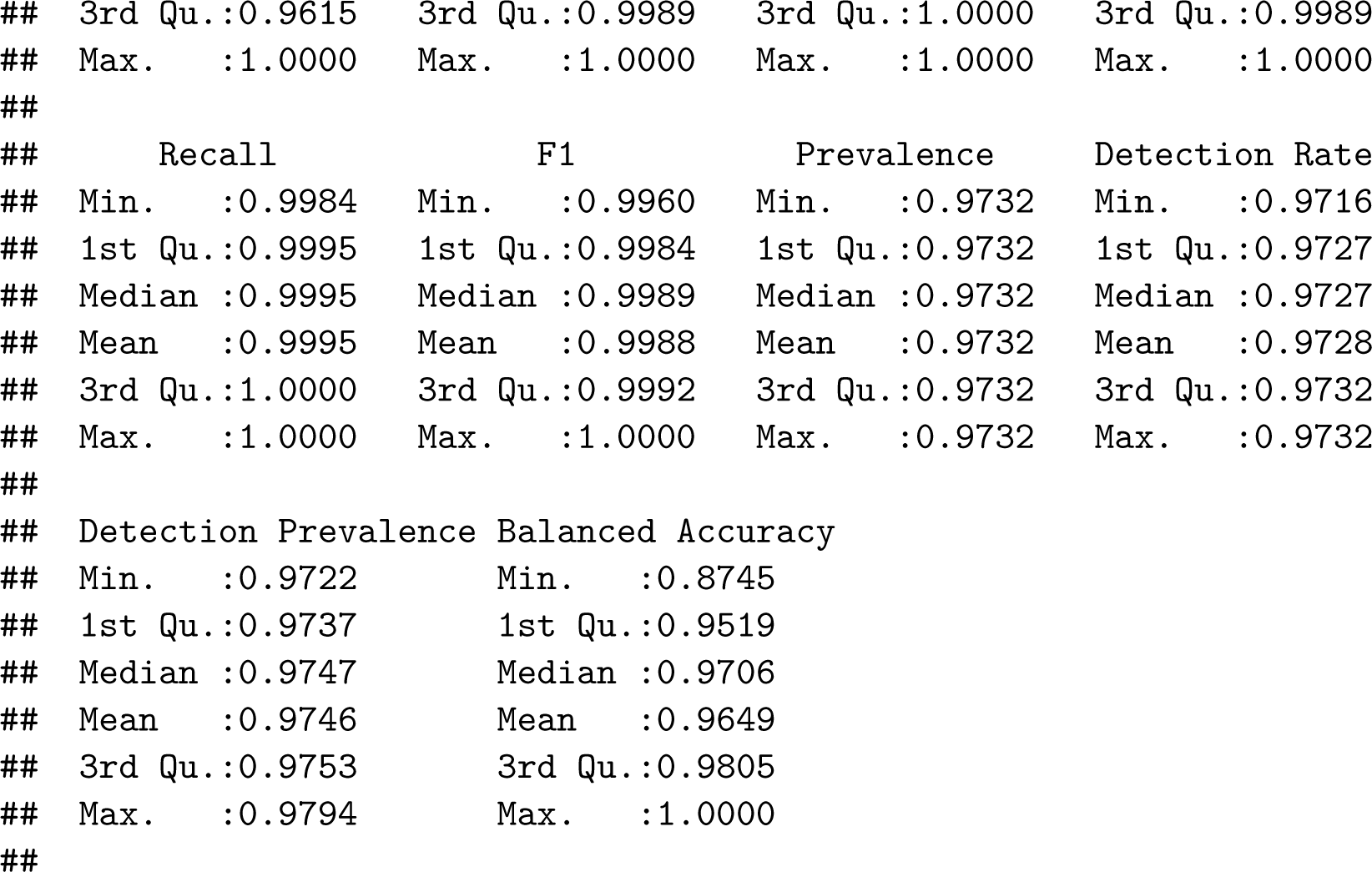

The performance of the PCA+QDA classifier using reference ‘Ref0’ is:

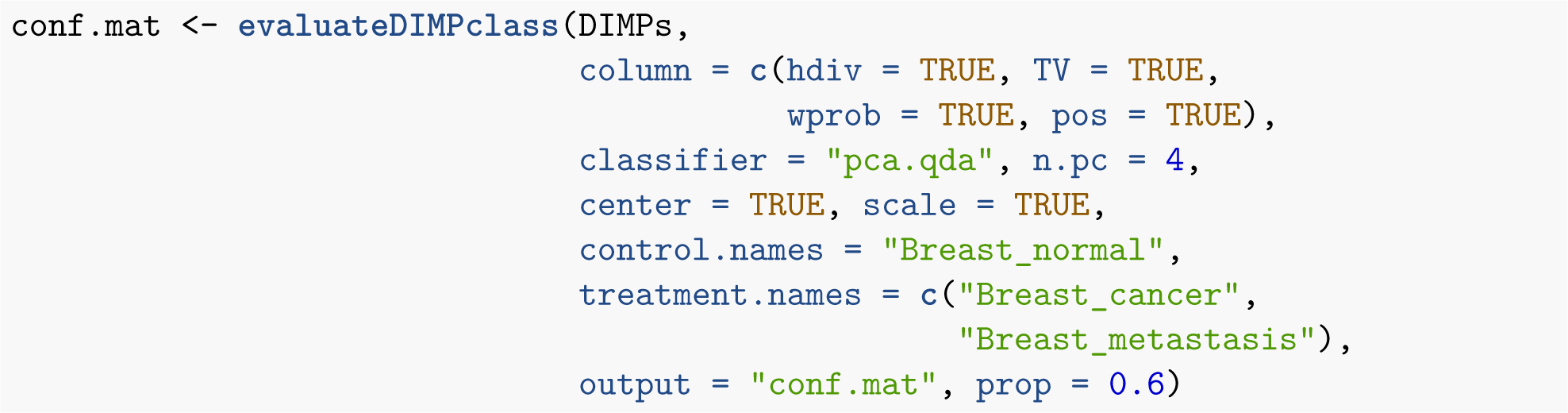

`## Model: treat ~ hdiv + TV + logP + pos`

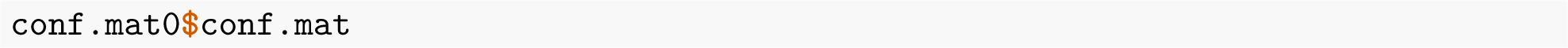

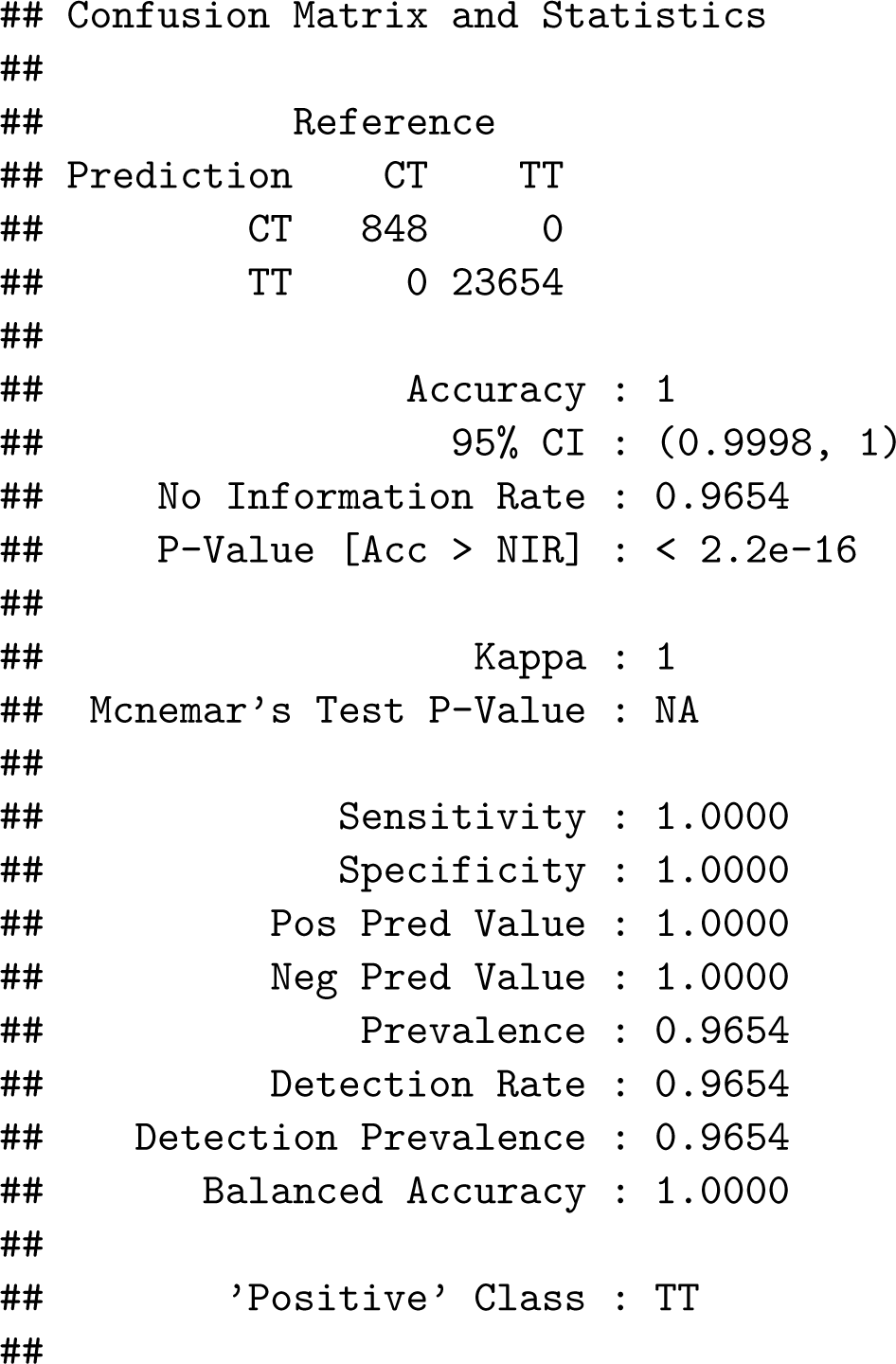

Monte Carlo (bootstrap) validation with 500 resamplings using reference ‘Ref0’ can be now performed:

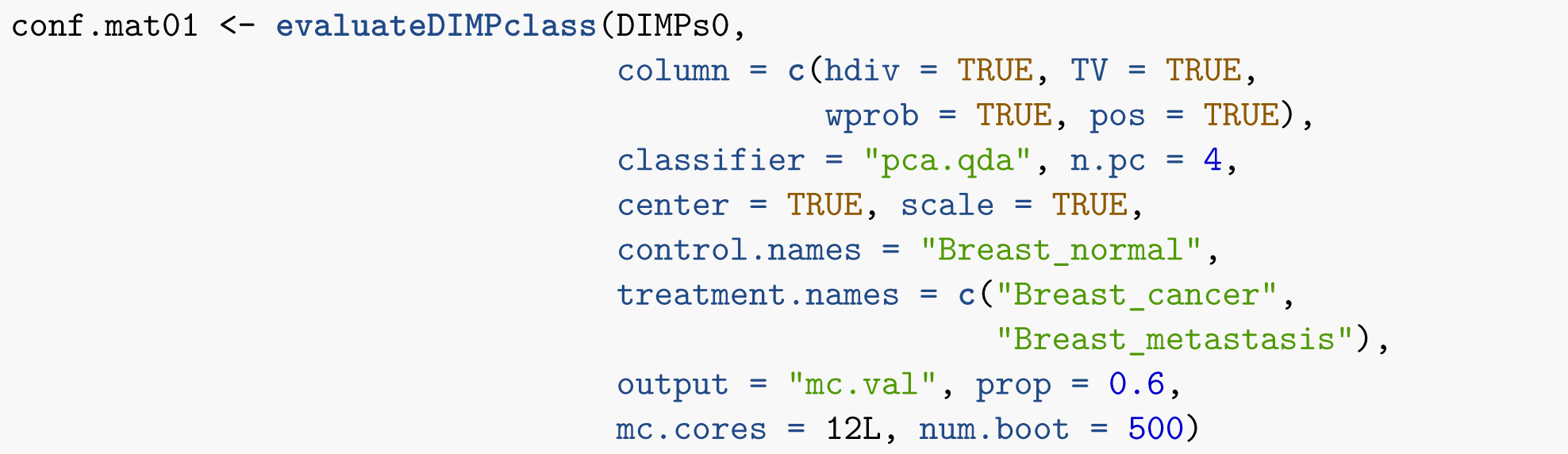

`## Model: treat ~ hdiv + TV + logP + pos`

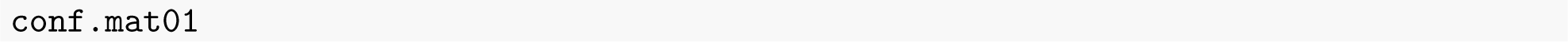

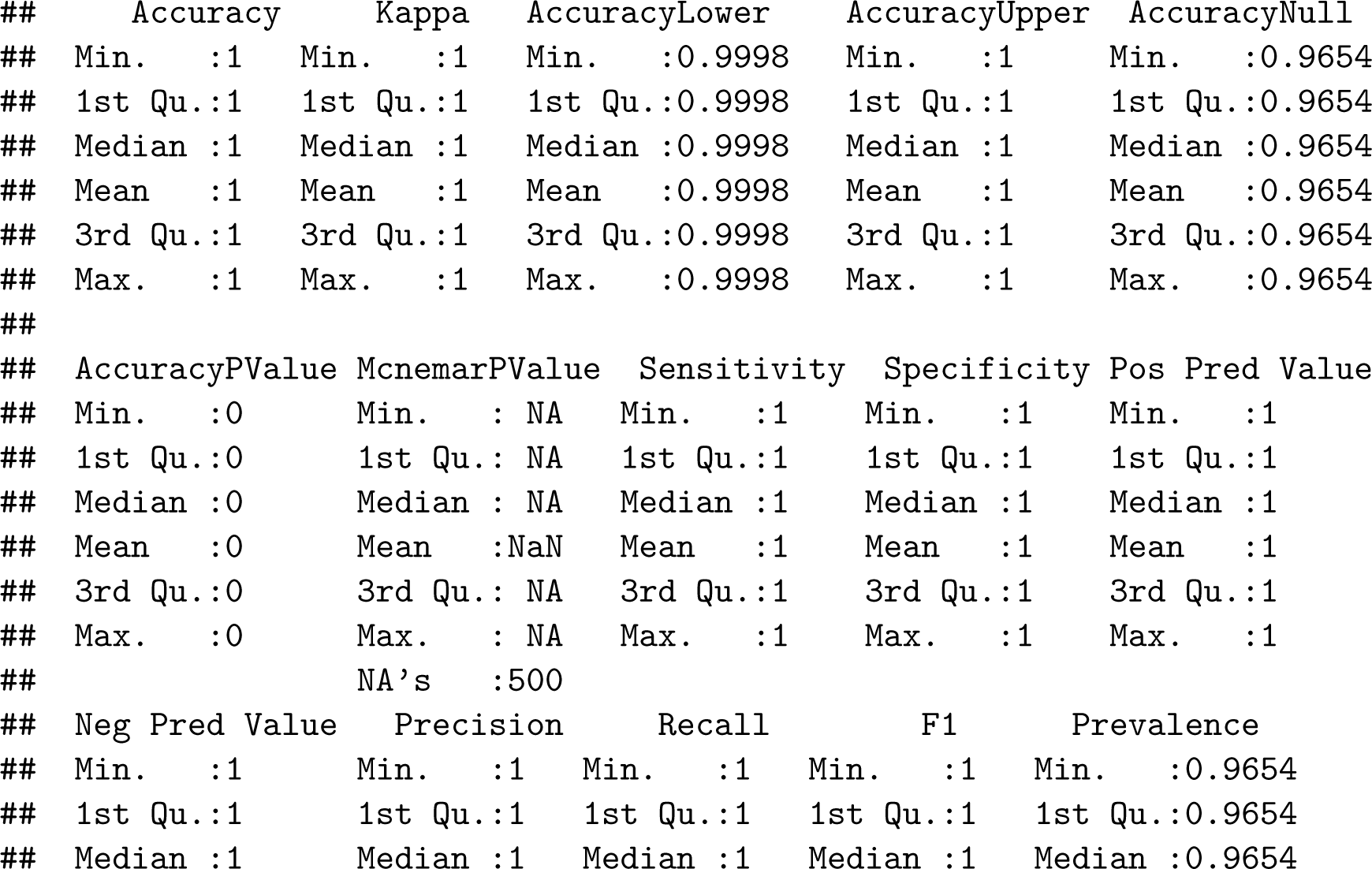
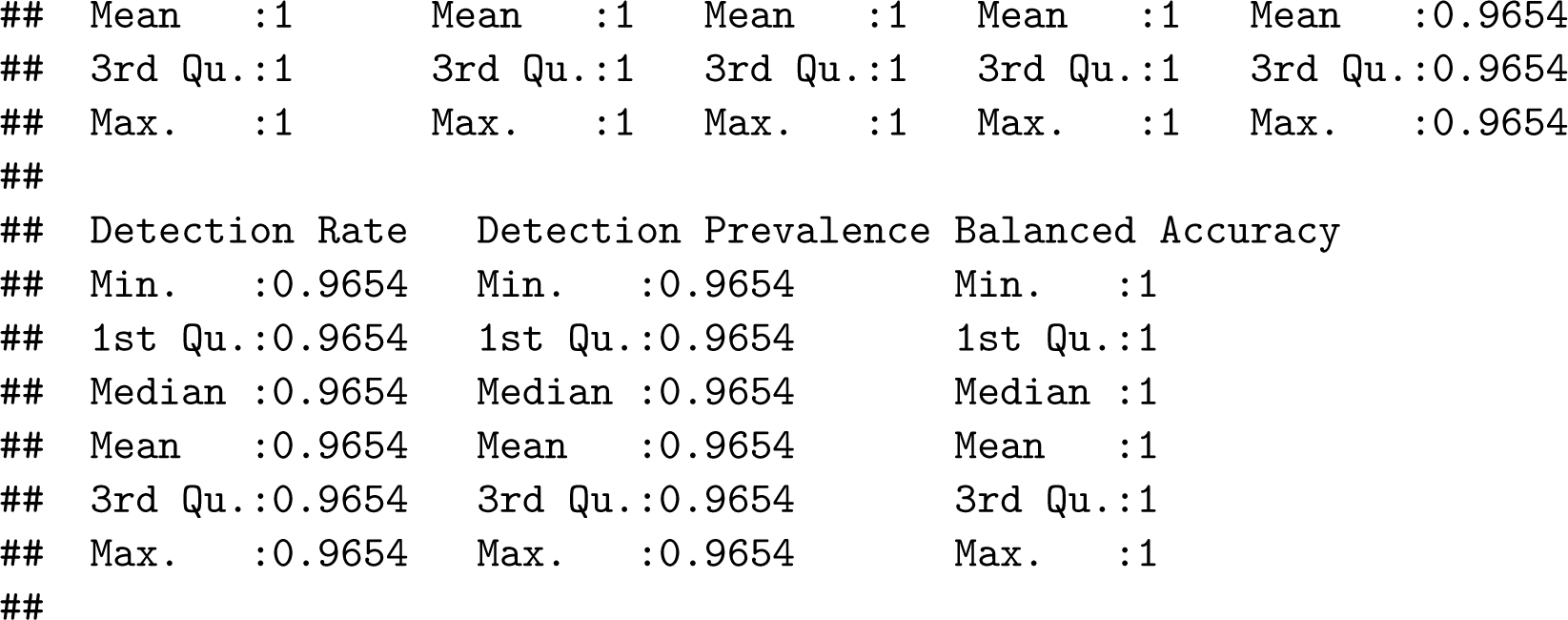

That is, with high accuracy level, DIMPs from control group can be discriminated from DIMPs found in cancer tissues.

## Acknowledgments

We thank Professor David J Miller for valuable conversations and suggestions on our mathematical modeling. # Funding The work was supported by funding from NSF-SBIR (2015-33610-23428-UNL) and the Bill and Melinda Gates Foundation (OPP1088661).

## Supplements

### S1. Troubleshooting installation on Ubuntu

Herein, a possible path to prevent potential issues originated during MethylIT installation on Ubuntu is given:

1. To update R:
  i. To added an R CRAN repository typing in the terminal: sudo echo “deb /bin/linux/ubuntu xenial/” | sudo tee -a /etc/apt/sources.list For example:

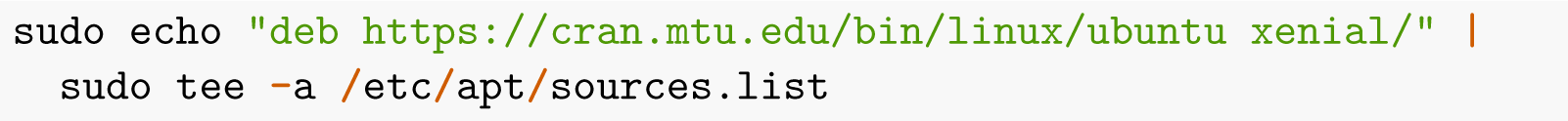
  ii. sudo apt update
  iii. sudo apt upgrade
2. Install Bioconductor: source(“https://bioconductor.org/biocLite.R”) biocLite()
3. Install Bioconductor packages: ‘GenomicFeatures’, ‘VariantAnnotation’, ‘ensembldb’, ‘GenomicRanges’, ‘BiocParallel’, ‘biovizBase’, ‘DESeq2’, and ‘genefilter’. Package ‘GenomicFeatures’ depends on the R package ‘RMySQL’, which is not in ’Bioconductor. To install “RMySQL” from CRAN you might require the ’installation of the library “libmysqlclient-dev”. If this is the case, ’then you can solve it by typing in the Ubuntu Teminal:

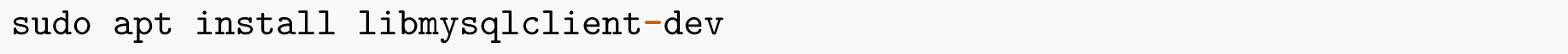 Next, in the R console:

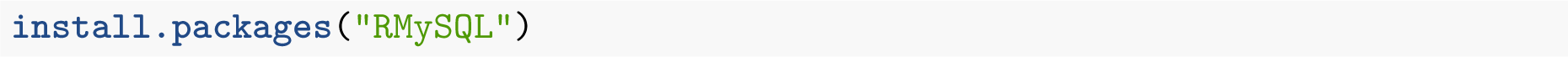
4. install.packages("devtools")
5. devtools::install_git("https://git.psu.edu/genomath/MethylIT")

### S2. Session Information

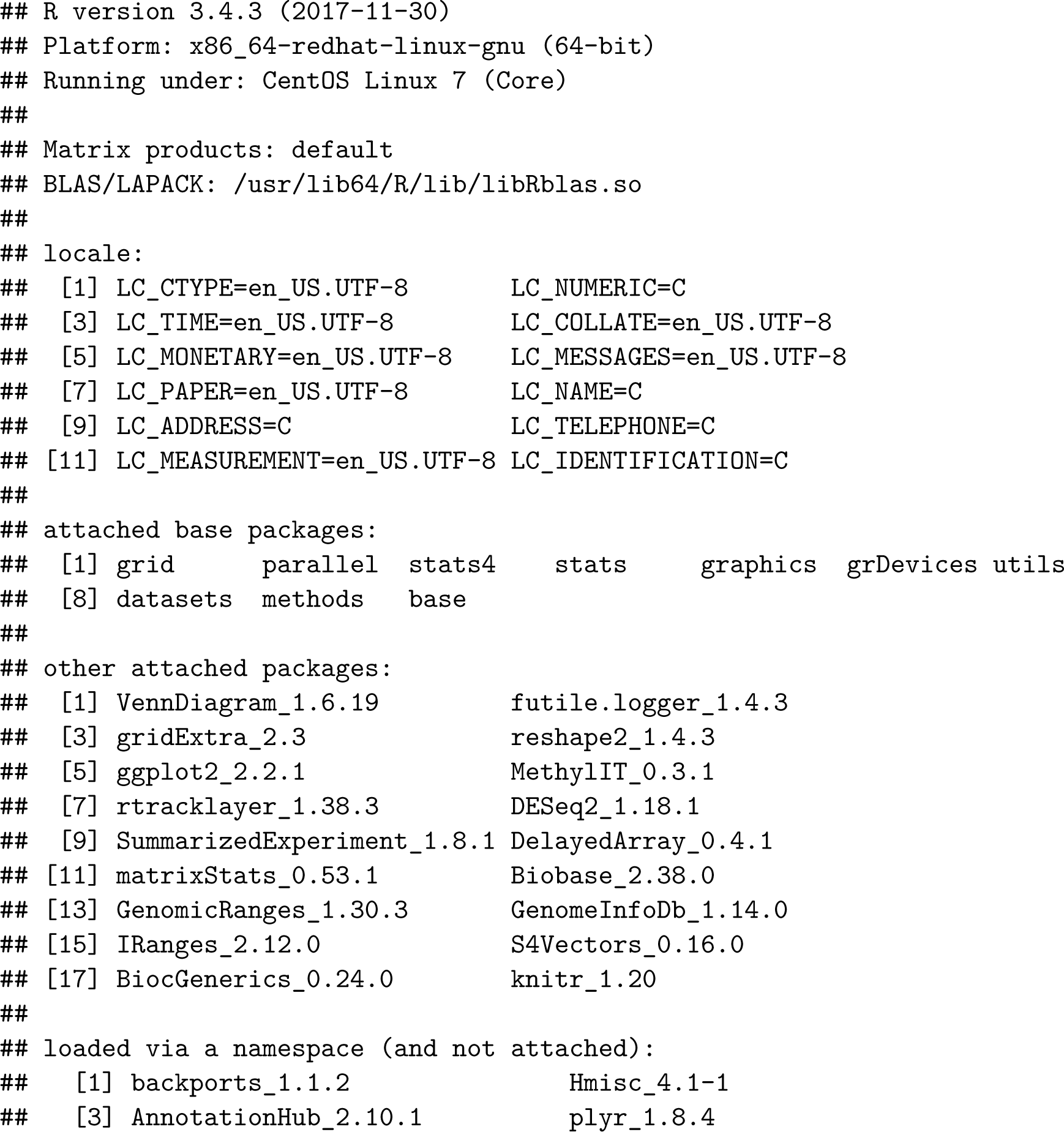
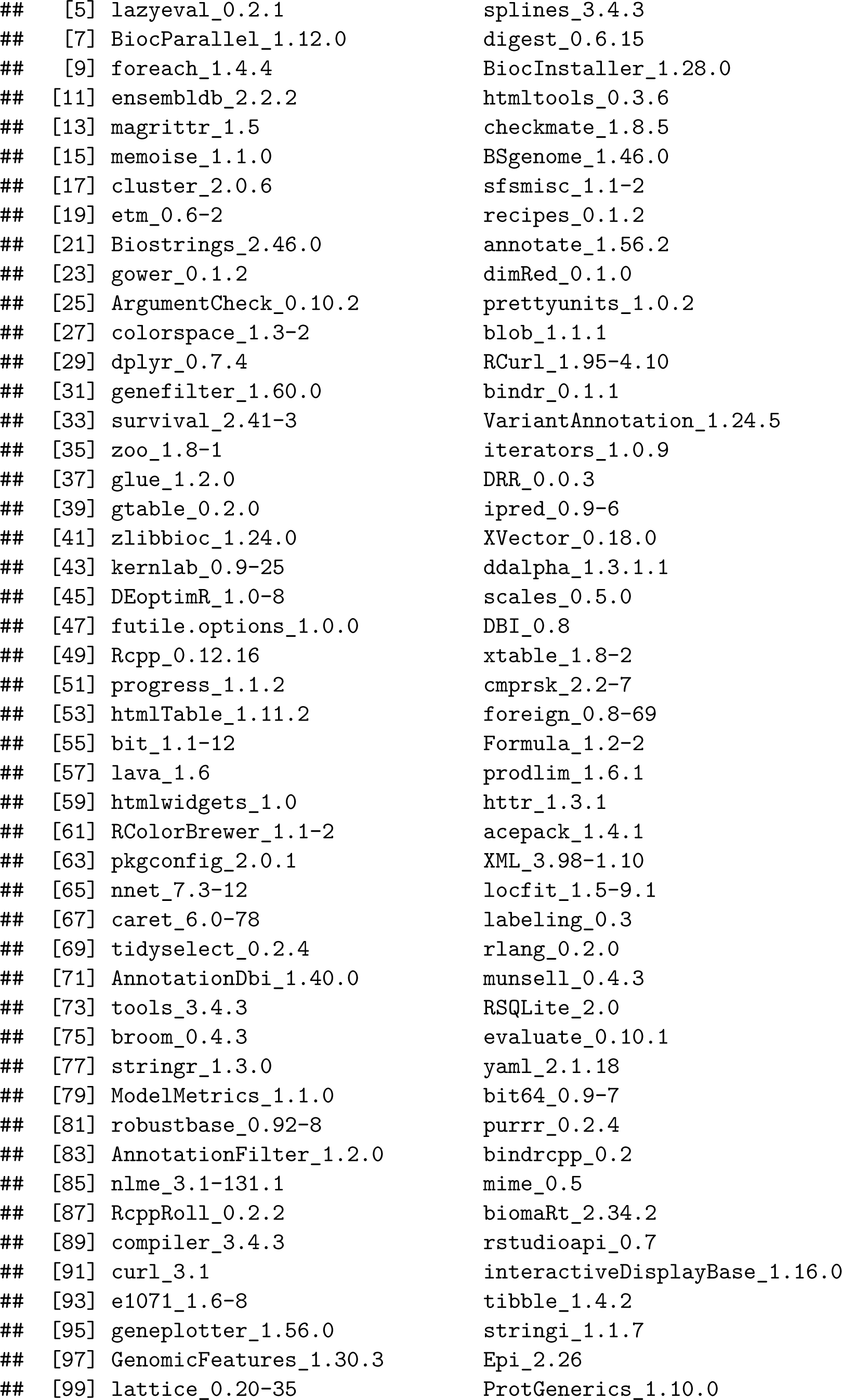
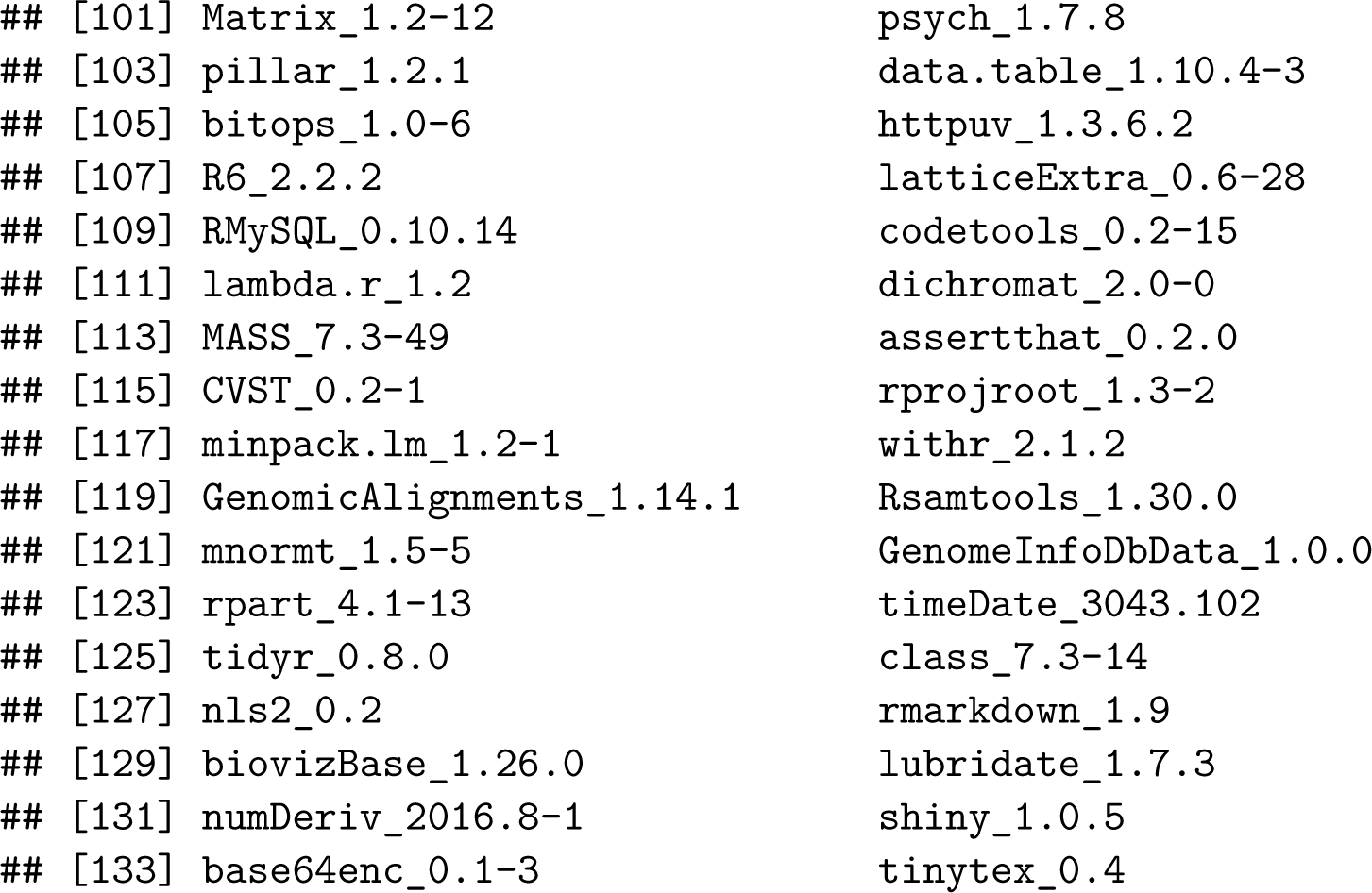

